# Molecular Evolution of *Pseudomonas syringae* Type III Secreted Effector Proteins

**DOI:** 10.1101/503219

**Authors:** Marcus M. Dillon, Renan N.D. Almeida, Bradley Laflamme, Alexandre Martel, Bevan S. Weir, Darrell Desveaux, David S. Guttman

## Abstract

Diverse Gram-negative pathogens like *Pseudomonas syringae* employ type III secreted effector (T3SE) proteins as primary virulence factors that combat host immunity and promote disease. T3SEs can also be recognized by plant hosts and activate an effector triggered immune (ETI) response that shifts the interaction back towards plant immunity. Consequently, T3SEs are pivotal in determining the virulence potential of individual *P. syringae* strains, and ultimately restrict *P. syringae* pathogens to a subset of potential hosts that are unable to recognize their repertoires of T3SEs. While a number of effector families are known to be present in the *P. syringae* species complex, one of the most persistent challenges has been documenting the complex variation in T3SE contents across a diverse collection of strains. Using the entire pan-genome of 494 *P. syringae* strains isolated from more than 100 hosts, we conducted a global analysis of all known and putative T3SEs. We identified a total of 14,613 T3SEs, 4,636 of which were unique at the amino acid level, and show that T3SE repertoires of different *P. syringae* strains vary dramatically, even among strains isolated from the same hosts. We also find that dramatic diversification has occurred within many T3SE families, and in many cases find strong signatures of positive selection. Furthermore, we identify multiple gene gain and loss events for several families, demonstrating an important role of horizontal gene transfer (HGT) in the evolution of *P. syringae* T3SEs. These analyses provide insight into the evolutionary history of *P. syringae* T3SEs as they co-evolve with the host immune system, and dramatically expand the database of *P. syringae* T3SEs alleles.

## INTRODUCTION

Over the past three decades, type III secreted effectors (T3SEs) have been recognized as primary mediators of many host-microbe interactions (Michiels and Cornelis, 1991;Salmond and Reeves, 1993;Hueck, 1998;Coburn et al., 2007;Deng et al., 2017;Hu et al., 2017;Rapisarda and Fronzes, 2018). These proteins are translocated directly from the pathogen cell into the host cytoplasm by the type III secretion system (T3SS), where they perform a variety of functions that generally promote virulence and suppress host immunity (Zhou and Chai, 2008;Cunnac et al., 2009;Oh et al., 2010;Buttner, 2016;Khan et al., 2018)}(Coburn et al., 2007). However, T3SEs can also be recognized by the host immune system, which allows the host to challenge the invading microbe. In plants, this immune response is called effector triggered immunity (ETI) (Jones and Dangl, 2006;Dodds and Rathjen, 2010;Khan et al., 2016). The interaction between pathogen T3SEs and the host immune system results in an evolutionary arms race, where pathogen T3SEs evolve to avoid detection while still maintaining their role in the virulence process, and the host immune system evolves to recognize the diversity of T3SEs and their actions, while maintaining a clear distinction between self and non-self to avoid autoimmune activation.

One of the best studied arsenals of T3SEs is carried by the plant pathogenic bacterium *Pseudomonas syringae* (Lindeberg et al., 2009;2012;Mansfield et al., 2012). *Pseudomonas syringae* is a highly diverse plant pathogenic species complex responsible for a wide-range of diseases on many agronomically important crop species (Mansfield et al., 2012). While the species as a whole has a very broad host range, individual strains can only cause disease on a small range of plant hosts (Sarkar et al., 2006;Lindeberg et al., 2009;Baltrus et al., 2017;Xin et al., 2018). A growing number of *P. syringae* strains have also recently been recovered from non-agricultural habitats, including wild plants, soil, lakes, rainwater, snow, and clouds (Morris et al., 2007;Morris et al., 2008;Clarke et al., 2010;Morris et al., 2013). This expanding collection of strains and the increased availability of comparative genomics data presents unique opportunities for obtaining insight into the determinants of host specificity in *P. syringae* (Baltrus et al., 2011;O’Brien et al., 2011;Baltrus et al., 2012;O’Brien et al., 2012;Dillon et al., 2017).

*Pseudomonas syringae* T3SEs have been the focus of both fundamental and applied plant pathology research for decades, going back to some of the early work on gene-for-gene resistance and avirulence proteins (Mukherjee et al., 1966;Staskawicz et al., 1984;Staskawicz et al., 1987;Keen and Staskawicz, 1988;Kobayashi et al., 1989;Keen, 1990;Jenner et al., 1991;Fillingham et al., 1992). Since then, over 1000 publications have focused on *P. syringae* T3SEs (Web of Science ["Pseudomonas syringae" AND (avirulence OR ("type III" AND effector))], October 2018), making it one of the most comprehensively studied T3SE systems. To date a total of 66 T3SE families and 764 T3SE alleles have been catalogued in the *Pseudomonas syringae* Genome Resources Homepage (https://pseudomonas-syringae.org). Many of these T3SE families are small, relatively conserved, and only distributed in a subset of *P. syringae* strains, while others are more diverse and distributed across the majority of sequenced *P. syringae* strains (Baltrus et al., 2011;O’Brien et al., 2011;Dillon et al., 2017). Given the irregular distribution of T3SEs among strains and their frequent association with mobile genetic elements, it has long been recognized that horizontal transfer plays an important role in the dissemination of T3SEs among strains (Kim and Alfano, 2002;Rohmer et al., 2004;Stavrinides and Guttman, 2004;Lovell et al., 2009;Godfrey et al., 2011;Lovell et al., 2011;Neale et al., 2016). Nucleotide composition and phylogenetic analyses of a subset of T3SEs identified eleven *P. syringae* T3SE families that were acquired by recent horizontal transfer events. However, the remaining thirteen families appeared to be ancestral and vertically inherited, suggesting that there is also an important role for pathoadaptation in the evolution of T3SEs (Rohmer et al., 2004;Stavrinides et al., 2006;O’Brien et al., 2011). While T3SE repertoires are thought to be key determinants of host specificity, strains with divergent repertoires are at times capable of causing disease on the same host (Almeida et al., 2009;O’Brien et al., 2011;Lindeberg et al., 2012;O’Brien et al., 2012), signifying that we have much to learn about the ways in which T3SEs contribute to *P. syringae* virulence.

Two major issues impact our current understanding of T3SE diversity in *P. syringae:* sampling bias and nomenclature. The current catalogue of T3SEs listed on the *Pseudomonas syringae* Genome Resources Homepage come from approximately 120 strains that represent only a subset of the phylogroups in the *P. syringae* species complex. Expanding this strain collection to a more diverse set will undoubtedly expand our understanding of diversity within T3SE families and reveal as-yet identified families. Another persistent issue in *P. syringae* comparative genomics has been the lack of benchmark standards for naming and assigning new T3SEs. While a standardized set of criteria for the identification and naming of *P. syringae* T3SEs have been published and broadly accepted (Lindeberg et al., 2005), the recommendation that new candidate T3SEs be subjected to rigorous phylogenetic analyses prior to family designation has not always been consistently employed. While this problem is not nearly as interesting from a biological perspective, it is very important operationally, since poor classification and naming practices can lead to substantial confusion and even spurious conclusions. Part of this issue stems from the fact that T3SEs are multidomain proteins that can share homology with multiple divergent T3SE families (Stavrinides et al., 2006;McCann and Guttman, 2008). At the time of their discovery, many families also had fewer than three T3SE alleles, making robust phylogenetic analyses impossible. Whatever the root cause, we are currently in a situation where many T3SEs are annotated without family assignment, some very similar T3SEs have been assigned to different T3SE families, and some highly divergent T3SEs are assigned to the same family based on short tracts of local similarity. This situation should be rectified in order to facilitate more comprehensive analyses of the role of T3SEs in the outcomes of host-pathogen interactions, particularly in light of the growing database of *P. syringae* genomics resources.

Here, we present an expanded catalogue of T3SEs in *P. syringae* and an updated phylogenetic analysis of the diversity within each T3SE family. We identified a total of 14,613 T3SEs from 494 *P. syringae* whole-genomes that include strains from 11 of the 13 *P. syringae* species complex phylogroups. These strains allowed us to redefine evolutionarily distinct family barriers for T3SEs, examine the distribution of each family across the *P. syringae* species complex, quantify the diversity within each T3SE family, and explore how T3SEs are inherited. By expanding and diversifying the database of confirmed and predicted *P. syringae* T3SEs and placing all alleles in an appropriate phylogenetic context, these analyses will ultimately enable more comprehensive studies of the roles of individual T3SEs in pathogenicity and allow us to more effectively explore the contribution of T3SEs to host specificity.

## METHODS

### Genome Sequencing, Assembly, and Gene Identification

Four hundred and ninety-four *P. syringae* species complex strains were analyzed (Supplemental Dataset S1), of which 102 assemblies were obtained from public sequence databases, including NCBI/GenBank, JGI/IMG-ER, and PATRIC (Markowitz et al., 2012;Wattam et al., 2014;Coordinators, 2018), and 392 strains were sequenced in house by the University of Toronto Center for the Analysis of Genome Evolution and Function (CAGEF). Two hundred and sixty-eight of these sequenced strains were provided by the International Collection of Microorganisms from Plants (ICMP). For the strains sequenced by CAGEF, DNA was isolated using the Gentra Puregene Yeast and Bacteria Kit (Qiagen, MD, USA), and purified DNA was then suspended in TE buffer and quantified with the Qubit dsDNA BR Assay Kit (ThermoFisher Scientific, NY, USA). Paired-end libraries were generated using the Illumina Nextera XT DNA Library Prep Kit following the manufacturer’s instructions (Illumina, CA, USA), with 96-way multiplexed indices and an average insert size of ~400 bps. All sequencing was performed on either the Illumina MISeq or GAIIx platform using V2 chemistry (300 cycles). Following sequencing, read quality was assessed with FastQC v.0.11.5 (Andrews, 2010) and low-quality bases and adapters were trimmed using Trimmomatic v0.36 (Bolger et al., 2014) (ILLUMINACLIP: NexteraPE-PE.fa, Maximum Mismatch = 2, PE Palindrome Match = 30, Adapter Read Match = 10, Maximum Adapter Length = 8; SLIDINGWINDOW: Window Size = 4, Average Quality = 5; MENLEN = 20). All genomes were then *de novo* assembled into contigs with CLC v4.2 (Mode = fb, Distance mode = ss, Minimum Read Distance = 180, Maximum Read Distance = 250, Minimum Contig Length = 1000). Raw reads were then re-mapped to the remaining contigs using samtools v1.5 with default settings to calculate the read coverage for each contig (Li and Durbin, 2009). Any contigs with a coverage depth of less than the average contig coverage by more than two standard deviations were filtered out of the assembly. Finally, gene prediction was performed on each genome using Prodigal v2.6.3 with default settings (Hyatt et al., 2010).

### Annotation and Family Delimitation of Type III Secreted Effectors

To characterize the effector repertoire of each of the 494 *P. syringae* species used in this study, we first downloaded all available *P. syringae* effector, helper, and chaperone sequences from three public databases: NCBI (18,120) (https://www.ncbi.nlm.nih.gov), Bean 2.0 (225) (Dong et al., 2015), and the *Pseudomonas syringae* Genome Resources Homepage (843) (https://pseudomonas-syringae.org). Using this database of 19,188 T3SE associated sequences in *P. syringae*, we then performed a BLASTP analysis to ensure that all sequences that we downloaded were assigned to appropriate families, which was essential given that many of the sequences downloaded from NCBI are ambiguously labelled as "type III effectors”, “type III helpers”, or “type III chaperones”. Any unassigned T3SE associated gene that had significant reciprocal blast hits (E < 1e-24) with an assigned T3SE associated gene was assigned to the corresponding family. This strict E-value cutoff was chosen to avoid incorrectly assigning families to sequences based on short-tracts of similarity that are common in the N-terminal region of T3SEs from different families (Stavrinides et al., 2006). Sequences that had reciprocal significant hits from multiple families were assigned to the family where they had more significant hits, which means that smaller families could be dissolved into a larger family if all sequences from the two families were sufficiently similar. However, this only occurred in one case, which resulted in all HopBB sequences being dissolved into the HopF family. In sum, our final seed database of *P. syringae* T3SEs contained a total of 7,974 effector alleles from 66 independent families, 1,585 discontinued effector alleles from 6 independent families, 2,230 helper alleles from 23 independent families, and 1,569 chaperones alleles from 10 independent families. Any sequences that were not able to be assigned to an appropriate T3SE family were discarded because of the possibility that these are not true T3SE associated genes.

Using the T3SE seed database, which contained a total of 7,974 effector alleles, we then annotated any predicted genes in each of the *P. syringae* genomes as a T3SE if the gene had a significant blast hit (E < 1e-24) in the T3SE seed database. This resulted in the annotation of 14,613 T3SEs across the 494 *P. syringae* strains. Family names were initially assigned to these T3SEs based on the name that had been assigned to the hit T3SE in the seed database. However, a meaningful comparative analysis of the distribution and evolution of the different T3SE families across the *P. syringae* species complex requires that we employ consistent definitions for delimiting each T3SE family. This has been historically problematic with *P. syringae* T3SEs because inconsistent criteria have been employed for assigning novel families. Therefore, we took all 14,613 T3SEs that were identified in this study and used an all-vs-all BLAST clustering approach to delimit them into new families with consistent criteria.

First, we blasted each T3SE amino acid sequence against a database of all 14,613 T3SEs and retained only hits that an E-value of less than 1e-24 and a length that covered at least 60% of the shorter sequence. Sequences that had multiple non-contiguous hits (i.e. high-scoring segment pairs) with an e-value less than 1e-24 whose cumulative lengths covered at least 60% of the shorter sequence were also retained. As was the case above, the strict e-value cutoff prevents us from assigning significant hits between T3SE sequences that only share strong local identity, which is most commonly seen in the N-terminal secretion signal. The 60% length cutoff prevents chimeric T3SEs from linking the two unrelated T3SE families that combined to form the chimera.

Second, a final list of all T3SE pairs that shared significant hits was gathered and T3SE sequences were collectively binned based on their similarity relationships. With this method, T3SE families were built based on all-by-all pairwise similarity between T3SEs rather than the similarity between individual T3SEs and an arbitrary seed T3SE or collection of centroid T3SEs, as is the case with some clustering methods. Significantly, our approach binned all significantly similar T3SE regardless of whether any two T3SEs were connected through direct or transitive similarity. For example, if T3SE sequence A was significantly similar to T3SE sequence B, and sequence B was significantly similar to sequence C, all three sequences would be binned together, regardless of whether there was significant similarity between sequence A and sequence C. This is important for appropriately clustering particularly diverse T3SE families, which may contain highly divergent alleles that have intermediate variants.

Finally, we assigned the same T3SE family designation to all T3SEs within each cluster based on the most commonly assigned T3SE family name that had initial been assigned to sequences within that cluster. In the majority of cases, all sequences in a single cluster had the same initially assigned T3SE family. However, for cases where there were multiple family names assigned to sequences within a single cluster, the lower Hop designation (ie. HopC < HopD) was assigned to all sequences in the cluster. Conversely, for cases where T3SEs that had initially been assigned the same family designation formed two separate clusters, T3SEs from the larger cluster were assigned the initial family name, and T3SEs from the smaller cluster(s) were assigned a novel family name, starting with HopBO, which is the first available Hop designation. Ultimately, this method allowed us to effectively delimit all T3SEs in this dataset into separate families with consistent definitions and performed considerably better at partitioning established T3SE families than standard orthology delimitation software like PorthoMCL (Tabari and Su, 2017) (Supplemental Dataset S2), likely because of the widespread presence of chimeric T3SEs in the *P. syringae* species complex.

In order to classify short chimeric relationships between families, as illustrated in Figure 2, we used a similar approach to the one outlined above. Specifically, we parsed our reciprocal BLASTP results to capture hits that occurred between alleles that had been assigned to different families. Here, we determined there to be a significant overlap between the alleles if there was an E-value < 1e-10, with no length limitation. These local relationships between some alleles in distinct families have no bearing on the evolutionary analyses performed in this study, but are highlighted in Figure 2, where the length of the alleles and their overlapping regions is proportional to the lengths of a pair of representative alleles from the two families.

### Phylogenetic Analyses

We generated three separate phylogenetic trees in this study to ask whether core-genome diversity, pan-genome content, or effector content could effectively sort *P. syringae* strains based on their host of isolation. For the core genome tree, we clustered all protein sequences from the 494 *P. syringae* genomes used in this study into ortholog families using PorthoMCL v3 with default settings (Tabari and Su, 2017). All ortholog families that were present in at least 95% of the *P. syringae* strains in our dataset were considered part of the soft-core genome and each of these families was independently aligned using MUSCLE v3.8.31 with default settings (Edgar, 2004). These alignments were then concatenated end-to-end using a custom python script and a maximum likelihood phylogenetic tree was constructed based on the concatenated alignment using FastTree v2.1.10 with default parameters (Price et al., 2010). For the pan-genome tree, we generated a binary presence-absence matrix for all ortholog families that were present in more than one *P. syringae* strain. This presence-absence matrix was used to compute a distance matrix in R v3.3.1 using the “dist” function with the Euclidean distance method. The phylogenetic tree was then constructed using the “hclust” function with the complete linkage hierarchical clustering method. We used the same approach to generate the effector content tree, except the input binary presence-absence matrix contained information on the 70 effector families rather than all ortholog families that made up the *P. syringae* pan-genome.

### Estimating Pairwise *Ka, Ks*, and *Ka/Ks*

Evolutionary rate parameters were calculated independently for each T3SE family. First, amino acid sequences were multiple aligned with MUSCLE v.3.8.31 using default settings (Edgar, 2004). Each multiple alignment was then reverse translated based on the corresponding nucleotide sequences using RevTrans v1.4 (Wernersson and Pedersen, 2003) and all pairwise *Ka* and *Ks* values were calculated for each family using the Nei-Gojobori Method, implemented by MEGA7-CC (Kumar et al., 2016). Output files were parsed using custom python scripts to convert the *Ka* and *Ks* matrices to stacked data frames with four columns: Sequence 1 Header, Sequence 2 Header, *Ka*, and *Ks*. The alignment-wide ratio of non-synonymous to synonymous substitutions (Ka/Ks) was then calculated for all T3SE pairs that had both a *Ka* and a *Ks* value greater than 0 in each family. For codon-level analysis of positive selection in each family, we used Fast Unconstrained Bayesian Approximation (FUBAR) to detect signatures of positive selection in all families that were present in at least five strains with default settings (Murrell et al., 2013).

For comparisons between T3SE family evolutionary rates and core genome evolutionary rates, we converted each individual core genome family alignment that was generated with MUSCLE to a nucleotide alignment with RevTrans, then concatenated these alignments end-to-end as described above. As was the case with each T3SE family, we then calculated *Ka* and *Ks* for all possible pairs of core genomes using the Nei Gojobori Method and parsed the output files into stacked data frames using our custom python script. The core genome data frame was then merged with each T3SE family data frame independently based on the genomes that the two T3SE sequences were from so that the evolutionary rates between these two T3SEs could be directly compared to the evolutionary rates of the corresponding core genomes.

### Gain-Loss Analysis

We used Gain Loss Mapping Engine (GLOOME) to estimate the number of gain and loss events that have occurred for each T3SE family over the course of the evolution of the *P. syringae* species complex (Cohen et al., 2010). The gain-loss analysis implemented by GLOOME integrates the presence-absence data for each gene family of interest across and the phylogenetic profile to estimate the posterior expectation of gain and loss across all branches. These events are then summed to calculate the total number of gene gain and loss events that have occurred for each family across the phylogenetic tree. We performed this analysis on each T3SE family using the mixture model with variable gain/loss ratio and a gamma rate distribution. The phylogenetic tree that used for this analysis was the concatenated core genome tree, which gives us the best estimation of the evolutionary relationships between strains, given the ample recombination known to occur within the *P. syringae* species complex (Dillon et al., 2017).

## RESULTS

In this study, we analyzed the type III effectorome of the *P. syringae* species complex using whole-genome assemblies from 494 strains representing 11 of the 13 established phylogroups and 72 distinct pathovars (Supplemental Dataset S1). These strains were isolated from 28 countries between 1935 and 2016, and include 62 *P. syringae* type and pathotype strains (Thakur et al., 2016). Although the majority of the strains were isolated from a diverse collection of more than 100 infected host species, we also included a number of strains isolated from environmental reservoirs, which have been dramatically under-sampled in *P. syringae* studies (Morris et al., 2007;Mohr et al., 2008;Clarke et al., 2010;Demba Diallo et al., 2012;Monteil et al., 2013;Morris et al., 2013;Monteil et al., 2016;Karasov et al., 2018). As per Dillon et al. (Dillon et al., 2017), we designate phylogroups 1, 2, 3, 4, 5, 6, and 10 as primary phylogroups and 7, 9, 11, and 13 as secondary phylogroups (we have no representatives from phylogroups 8 or 12, although presumably they would also be secondary phylogroups) (Berge et al., 2014). The primary phylogroups are phylogenetically quite distinct from the secondary phylogroups and include all of the well-studied *P. syringae* strains. Nearly all of the primary phylogroup strains carry a canonical *P. syringae* type III secretion system and were isolated from plant hosts. In contrast, many of the strains in the secondary phylogroups do not carry a canonical *P. syringae* type III secretion system and were isolated from environmental reservoirs (e.g. soil or water).

All of the *P. syringae* genome assemblies used in this study were downloaded directly from NCBI or generated in-house by the University of Toronto Centre for the Analysis of Genome Evolution & Function using paired-end data from the Illumina GAIIx or the Illumina MiSeq platform. There was some variation in the genome sizes, contig numbers, and N50s among strains due to the fact that the majority of the genomes are *de novo* assemblies in draft format (Figure S1); however, the number of coding sequences identified in each strain were largely consistent with the six finished (closed and complete) genome assemblies in our dataset. Given the large size of the *P. syringae* pan-genome, the fact that some strains have acquired large plasmids, and the relatively high frequency of horizontal gene transfer in the *P. syringae* species complex (Baltrus et al., 2011;Dillon et al., 2017), we expect there to be some variation in genome size and coding content of different strains.

### Distribution of type III secreted effectors in the *P. syringae* species complex

To explore the distribution of T3SEs across the *P. syringae* species complex, we first identified all putative T3SEs present in each of our 494 genome assemblies using a blastp analysis (Altschul et al., 1997), where all protein sequences from each *P. syringae* genome were queried against a database of known *P. syringae* T3SEs obtained from the *Pseudomonas syringae* Genome Resource Database (https://pseudomonas-syringae.org), the Bean 2.0 T3SE Database (http://systbio.cau.edu.cn/bean), and the NCBI Protein Database (https://www.ncbi.nlm.nih.gov). In sum, we identified a total of 14,613 confirmed and putative T3SEs, 4,636 of which were unique at the amino acid level, and 5,127 of which were unique at the nucleotide level. Individual *P. syringae* strains in the dataset harbored between one and 53 putative T3SEs, with a mean of 29.58 ± 10.13 (stddev), highlighting considerable variation in both the composition and size of each strain’s suite of T3SEs (Figure 1). As expected, primary phylogroup strains tended to harbor substantially more T3SEs than secondary phylogroups strains (30.55 ± 8.97 vs. 3.89 ± 1.64, respectively), which frequently do not contain a canonical T3SS (Dillon et al., 2017). However, a subset of strains from phylogroups 2 and 3, and all strains from phylogroup 10 harbored fewer than 10 T3SEs, more closely mirroring secondary phylogroup strains in their T3SE content. The extensive T3SE repertoires found in most primary phylogroup strains supports the idea that these effectors play an important role in the ecological interactions of the majority of strains in this species complex.

**Figure 1:**
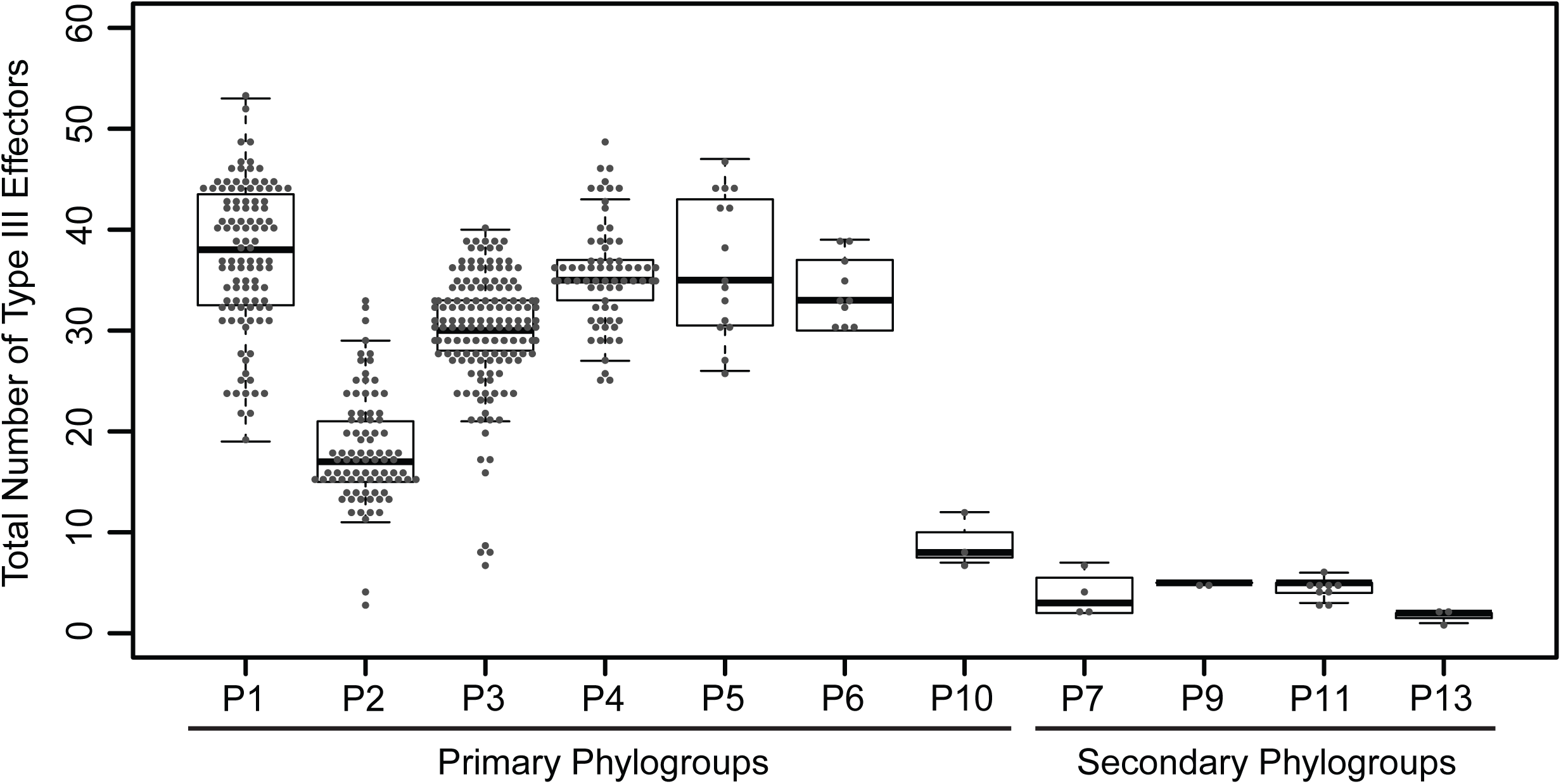
Total number of coding T3SEs in each *P. syringae* strain, sorted by phylogroup. Closed circles represent the number of effectors in each strain, boxes show the first quartile effector count, median effector count, and third quartile effector count for the whole phylogroup, and whiskers extend to the highest and lowest effector counts in the phylogroup that are not identified as outliers (>1.5 times the interquartile range).

Objective criteria are required for partitioning and classifying T3SEs prior to any study of their distribution and evolution. In 2005, an effort was made to unify the disparate classification and naming conventions applied to *P. syringae* T3SEs (Lindeberg et al., 2005). While this effort was very successful overall, the criteria have not been universally or consistently applied, resulting in some problematic families. For example, the HopK and AvrRps4 families are homologous over the majority of their protein sequences, but are assigned to distinct families, while the HopX family contains highly divergent subfamilies that only share short tracts of local similarity.

We reassessed the relationship between all 14,613 T3SEs using a formalized protocol in order to objectively delimit families and clarify the current classification. While the selection of the specific delimiting criteria is arbitrary and open to debate, we have elected to use a well-established protocol with fairly conservative thresholds. We identified shared similarity using a BLASTP-based pairwise reciprocal best hit approach (Altschul et al., 1997;Eisen, 2000;Daubin et al., 2002), with a stringent Expect-value acceptance threshold of E<1e-24 and a length coverage cutoff of ≥60% of the shorter sequence (regardless of whether it is query or subject). It should be noted that since this approach uses BLAST it requires only local similarity between family members. Nevertheless, our stringent E-value and coverage thresholds select for matches that share more extensive similarity than would typically be observed when proteins only share a single domain. We feel that these criteria provide a reasonable compromise between very relaxed local similarity criteria (using default BLAST parameters) and very conservative global similarity criteria. All T3SEs that exceeded our acceptance thresholds were sorted into family bins. T3SEs in each bin can therefore be either connected through direct similarity or transitive similarity. Finally, we assigned a name to all T3SEs in each bin based on the most common effector family name in that bin.

Our analysis identified 70 T3SE families and sorted T3SEs into their historical families in the majority of cases. However, there were some exceptions, including merging existing effector families that shared significant local similarity (Table 1), and creating some new, putative families that were generated from T3SEs originally assigned to existing families, but which did not pass our local similarity thresholds (Table 2). A number of these new families only contain a single allele, so it is likely that they are recent pseudogenes still annotated as coding sequences by Prodigal. Finally, in two cases, a subset of alleles from one T3SE family were assigned to a different family due to the extent of shared local similarity. This included the assignment of all originally designated HopS1 subfamily alleles to HopO, and the assignment of all originally designated HopX3 alleles to HopF.

**Table 1:** T3SE Families Merged into a New Family

| Families to Merge | New Family <sup>1</sup> |
| --- | --- |
| HopAB + HopAY | HopAB |
| HopAT + HopAV | HopAT |
| HopB + HopAC | HopB |
| HopAO + HopD | HopD |
| HopF + HopBB | HopF |
| HopK + AvrRps4 | HopK |
| HopW + HopAE | HopW |

The new name was assigned based on the first assigned Hop designation.

**Table 2:** New T3SE Families

| Old Name | New Family |
| --- | --- |
| HopX2 | HopBO |
| HopZ3 | HopBP |
| HopH3 | HopBQ |
| HopBN1 | HopBR |
| HopAV1 | HopBS |
| HopAB2 | HopBT <sup>1</sup> |
| HopAB2 | HopBU <sup>1</sup> |
| HopAJ2 | HopBV <sup>1</sup> |
| HopBH1 | HopBW <sup>1</sup> |
| HopL1 | HopBX <sup>1</sup> |

<sup>1</sup> These new families only contain a single allele

It is important to emphasize that the new criteria do not bin T3SEs that share less than 60% similarity across the shortest sequence. This was done to prevent families from being combined due to short chimeric relationships between a subset of the alleles in distinct families (Stavrinides et al., 2006). These relationships could be recognized as super-families, although the reticulated nature of these relationships makes this unwieldly. We list families that share these short regions of similarity in Figure 2, although it is important to recognize that some of these chimeric relationships are only displayed by a subset of alleles in each family. While we acknowledge that some of the new T3SE family boundaries may cause concern due to conflicts with historical naming, we feel it is essential to use unambiguous and consistent criteria for family delimitation.

**Figure 2:**
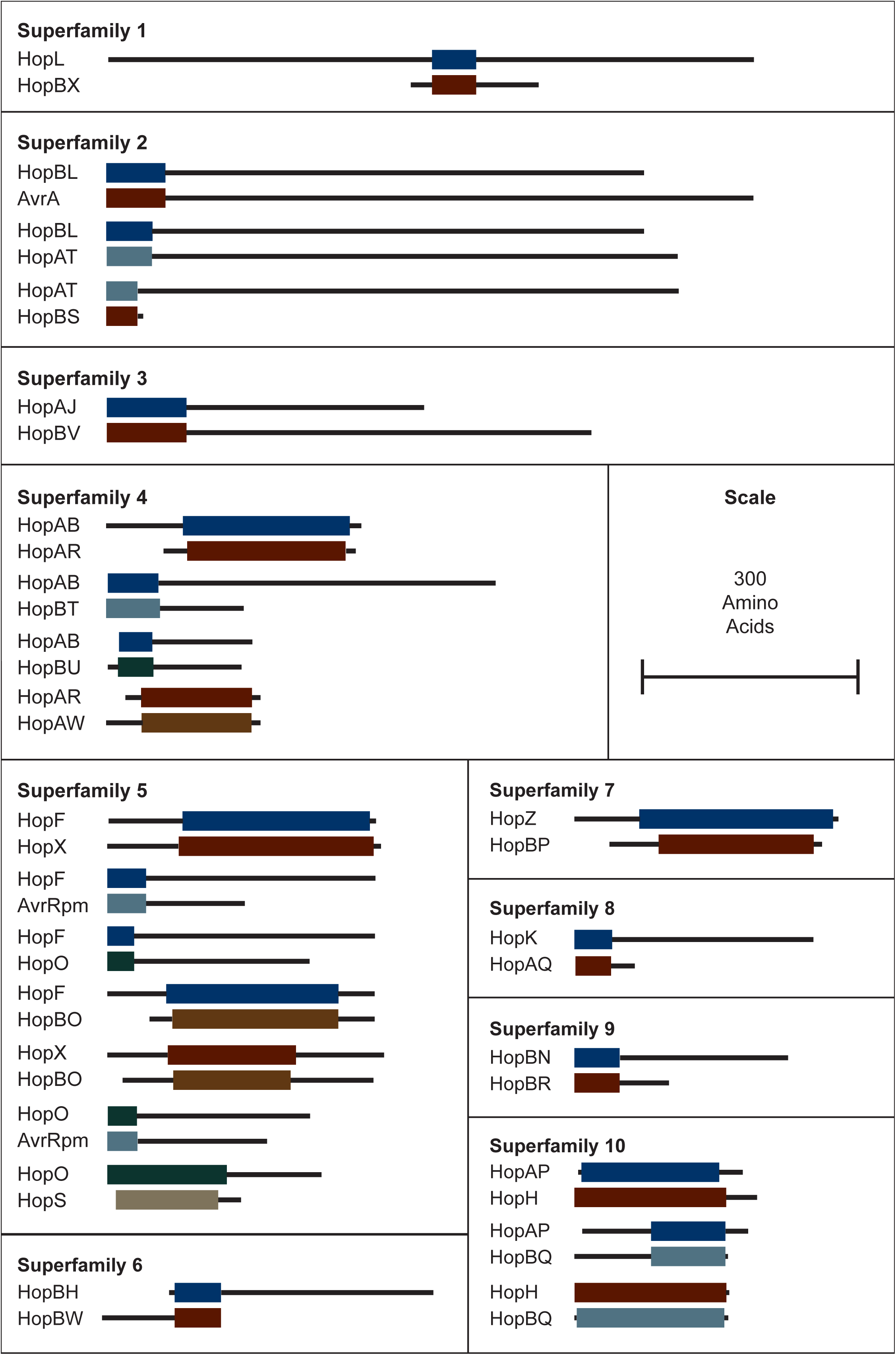
Interfamily blast hits (E < 1e-10) that did not pass our e-value and/or length cut-offs for combining T3SEs into families. Each superfamily represents a cluster of families that have some overlapping sequence. Coloured blocks represent the regions of the representative sequence pairs that are homologous, where the length of the blocks is proportional to the length of the homologous sequence. Black lines represent the remainder of each representative sequence that is not homologous, where the length of the lines is proportional to the length of the 5’ and 3’ non-homologous regions. Not all families within a superfamily need to contain a significant blast hit with all other families in the superfamily because they can be homologous to the same intermediate sequence in different regions.

The distribution of each of these 70 T3SE families across the *P. syringae* species complex reveals that the majority of families are present in only a small subset of *P. syringae* strains, typically from a few primary phylogroups (Figure 3; Figure S2). Among T3SE effector families, only AvrE, HopB, HopM, and HopAA are considered part of the soft-core genome of *P. syringae* (present in > 95% of strains). Interestingly, three of these core families, AvrE, HopM, and HopAA are part of the conserved effector locus (CEL), a well characterized and evolutionarily conserved sequence region that is present in most *P. syringae* strains (Alfano et al., 2000;Dillon et al., 2017). However, the fourth effector from the CEL, HopN, is only present in 14.98% of strains, all of which are from phylogroup 1. While the remainder of T3SE families are also mostly present in a small subset of strains, there is a wide distribution in the number of strains harboring individual T3SE families, further highlighting the dramatic variation in T3SE content across *P. syringae* strains (Figure S3).

**Figure 3:**
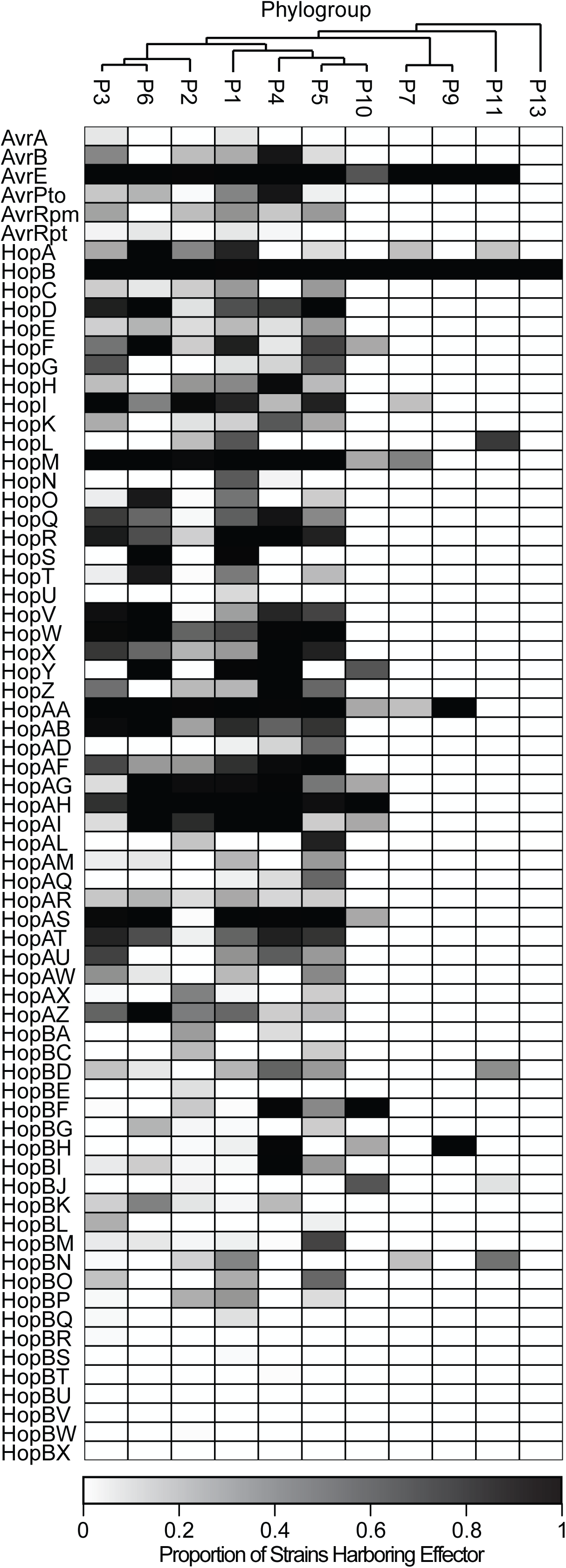
Heat map demonstrating the proportion of strains in each phylogroup that harbor each of the T3SE families. Only four T3SE families, AvrE, HopB, HopM, and HopAA are considered part of the soft-core *P. syringae* complex genome (present in > 95% of strains). Other T3SE families are mostly sparsely distributed across the *P. syringae* species complex, with several families only being present in a few phylogroups.

Following family and strain T3SE classification, we also performed hierarchical clustering using the T3SE content of each strain to determine if T3SE profiles are a good predictor of host specificity. We previously reported that in *P. syringae*, neither the core genome or gene content phylogenetic trees correlate well with the hosts from which the strains were isolated (Dillon et al., 2017). This remains true in this study, where we’ve updated the core and pan-genome analyses with an expanded set of strains (Figure S4; Figure S5). The T3SE content tree is not as well resolved due to the smaller number of phylogenetically informative signals in the T3SE dataset. However, we were able to largely recapitulate the established *P. syringae* phylogroups with this analysis, suggesting that more closely related strains do tend to have more similar T3SE repertoires (Figure S6). We also see that the phylogroup 2, phylogroup 3, and phylogroup 10 strains that have smaller T3SE repertoires than other primary phylogroups, cluster more closely with secondary phylogroup strains in the effectorome tree. However, as was the case in the core genome and gene content trees, hierarchical clustering based on effector content did not effectively separate strains based on their host of isolation. We therefore conclude that overall T3SE content is not a good predictor of host specificity.

### Diversification of type III secreted effectors in the *P. syringae* species complex

Substantial genetic and functional diversity has been shown to exist within individual T3SE families (Lewis et al., 2014;Dillon et al., 2017). While some T3SE families are relatively small, restricted to only a subset of *P. syringae* strains, and present in only a single copy in each strain, others are found in nearly all strains, and often appear in multiple copies within a single genome (Figure 4). Many of the largest families, including those that are part of the core genome (AvrE, HopB, HopM, and HopAA), are among those that are often present in multiple copies. However, we also found that some families that are present in less than half of *P. syringae* strains (e.g. HopF, HopO, HopZ, and HopBL) frequently appear in multiple copies. The average copy number of individual T3SEs per strain across all families is 1.30, while some families are present in copy numbers as high as six.

**Figure 4:**
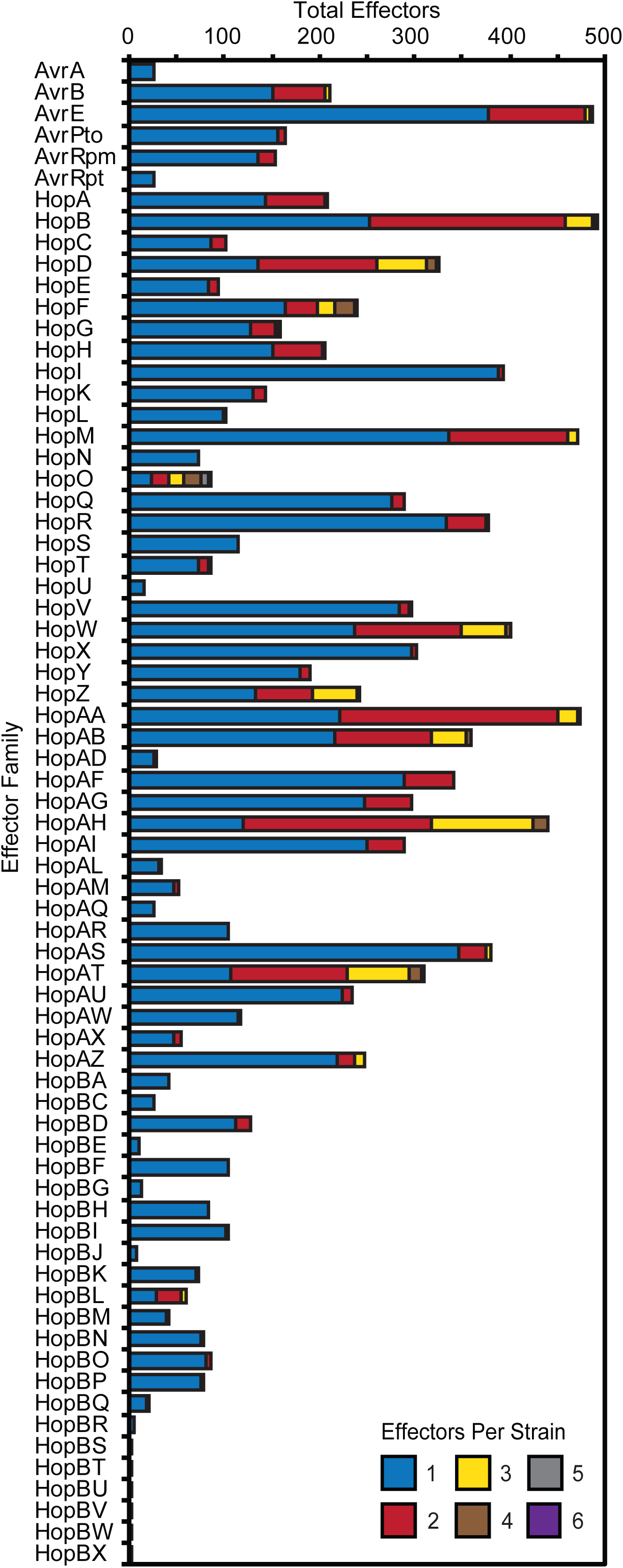
Total number of *P. syringae* strains harboring an allele from each T3SE family. Colour categories denote the copy number of each effector family in the corresponding strains. While the majority of families are mostly present in a single copy, some of the more broadly distributed families have higher copy numbers in a subset of *P. syringae* genomes.

To quantify the extent of genetic diversification within each T3SE family, we aligned the amino acid sequences of all members from each family with MUSCLE, then reverse translated these amino acid alignments and calculated all pairwise non-synonymous (*Ka*) and synonymous (*Ks*) substitution rates for all pairs of alleles within each family. There was a broad range of pairwise substitution rates in the majority of T3SE families, which is expected given the range of divergence times in the core-genomes of strains from different *P. syringae* phylogroups (Dillon et al., 2017). The three families with the highest non-synonymous substitution rates were HopF, HopAB, and HopAT (Figure 5A), which all have an average *Ka* greater than 0.5. These families also tended to have relatively high synonymous substitution rates, but several other families also have *Ks* values that are greater than 1.0 (Figure 5B).

**Figure 5:**
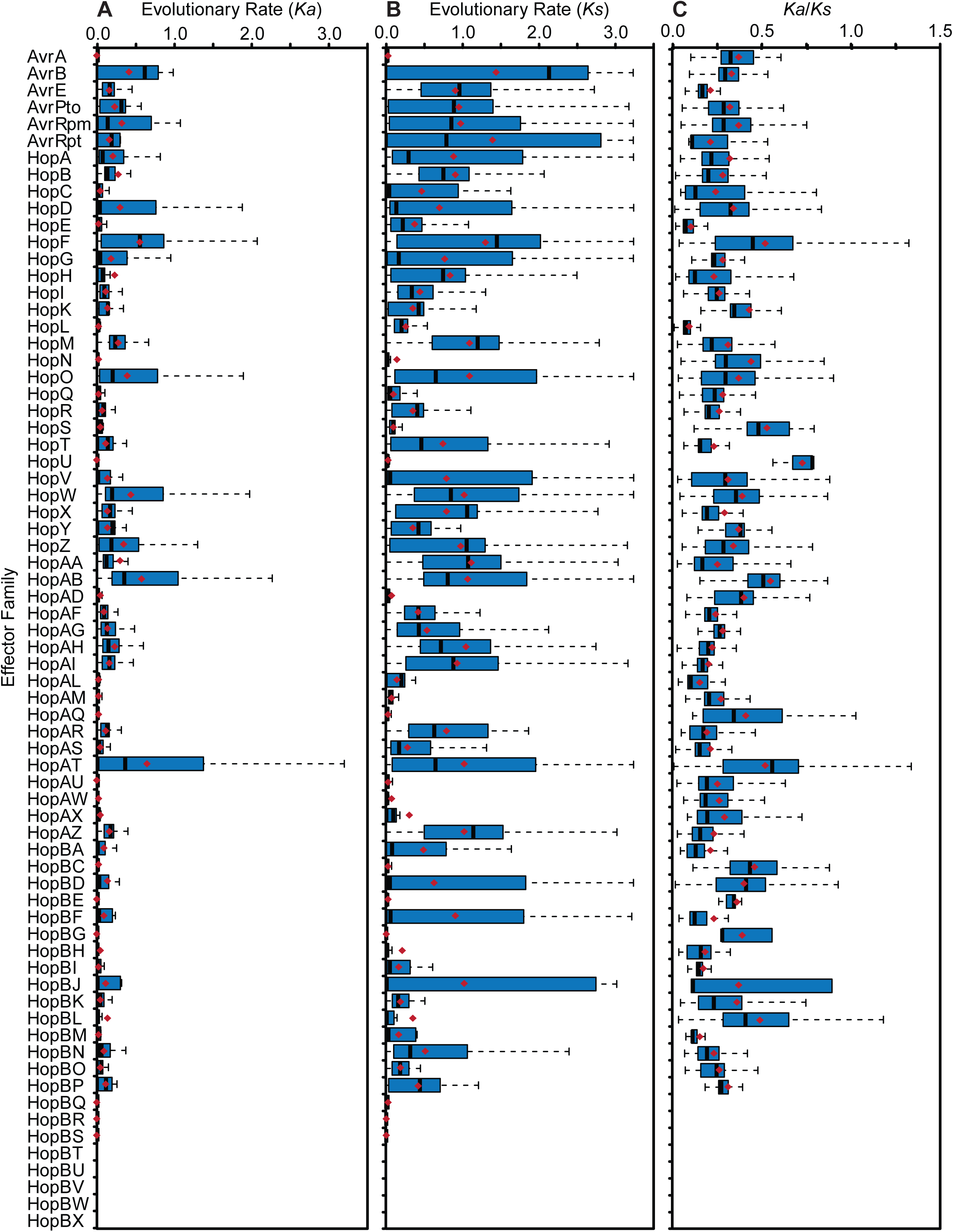
Non-synonymous substitution rate (*Ka*), synonymous substitution rates (*Ks*), and *Ka/Ks* ratio for each T3SE family. All alleles in each family were aligned using MUSCLE v. 3.8 and all pairwise *Ka* and *Ks* values within each family were calculated using MEGA7 with the Nei-Gojobori Method. Boxes show the first quartile substitution rates, median substitution rates, and third quartile substitution rates for each family, and whiskers extend to the highest and lowest substitution rates in the family that are not identified as outliers (>1.5 times the interquartile range). Average pairwise *Ka, Ks*, and *Ka/Ks* values for each family are denoted by red X’s.

While some pairwise comparisons of effector alleles did yield a *Ka/Ks* ratio greater than 1, the predominance of purifying selection operating in the conserved domains of these families likely overwhelms signals of positive selection at individual sites. Indeed, the average global pairwise *Ka/Ks* values were less than 1 for all T3SE families (Figure 5C). Therefore, we also analyzed the *Ka* and *Ks* on a per codon basis using FUBAR to search for site-specific signals of positive selection in each family (Bayes Empirical Bayes P-Value ≥ 0.9; *Ka/Ks >* 1) (Murrell et al., 2013). We find that 37 out of the 64 (57.81%) T3SE families with at least five alleles have at least one positively selected site. The number of positively selected sites in these families was relatively low, ranging from 1 to 17, with the percentage of positively selected sites in a single family never rising above 2.29% (Table 3). By comparison, we found that only 3,888/17,807 (21.83%) ortholog families from the pangenome of *P. syringae* that were present in at least five strains demonstrated signatures of positive selection at one or more sites (Dillon et al., 2017), suggesting that T3SE families experience extremely high rates of positive selection.

**Table 3:**
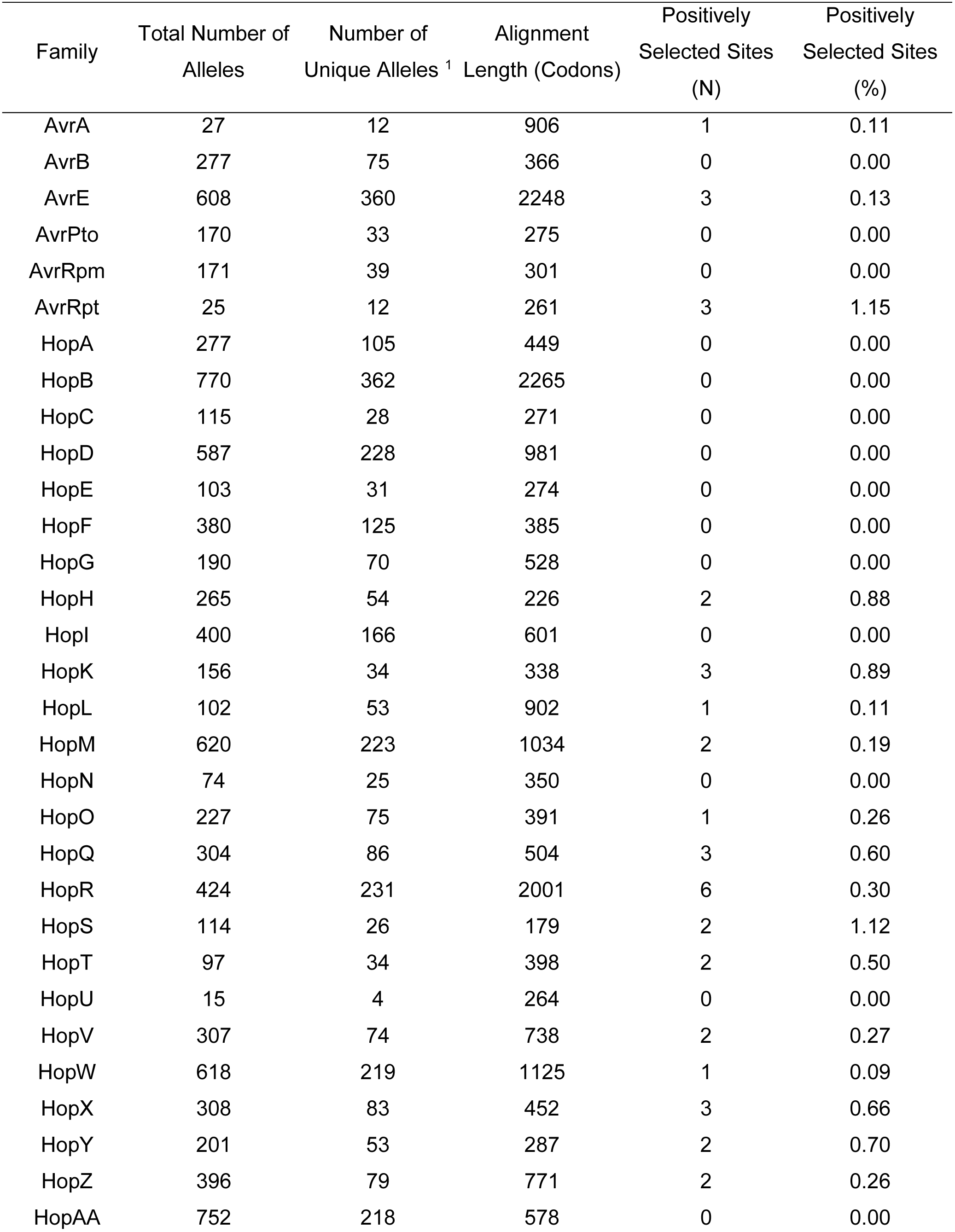
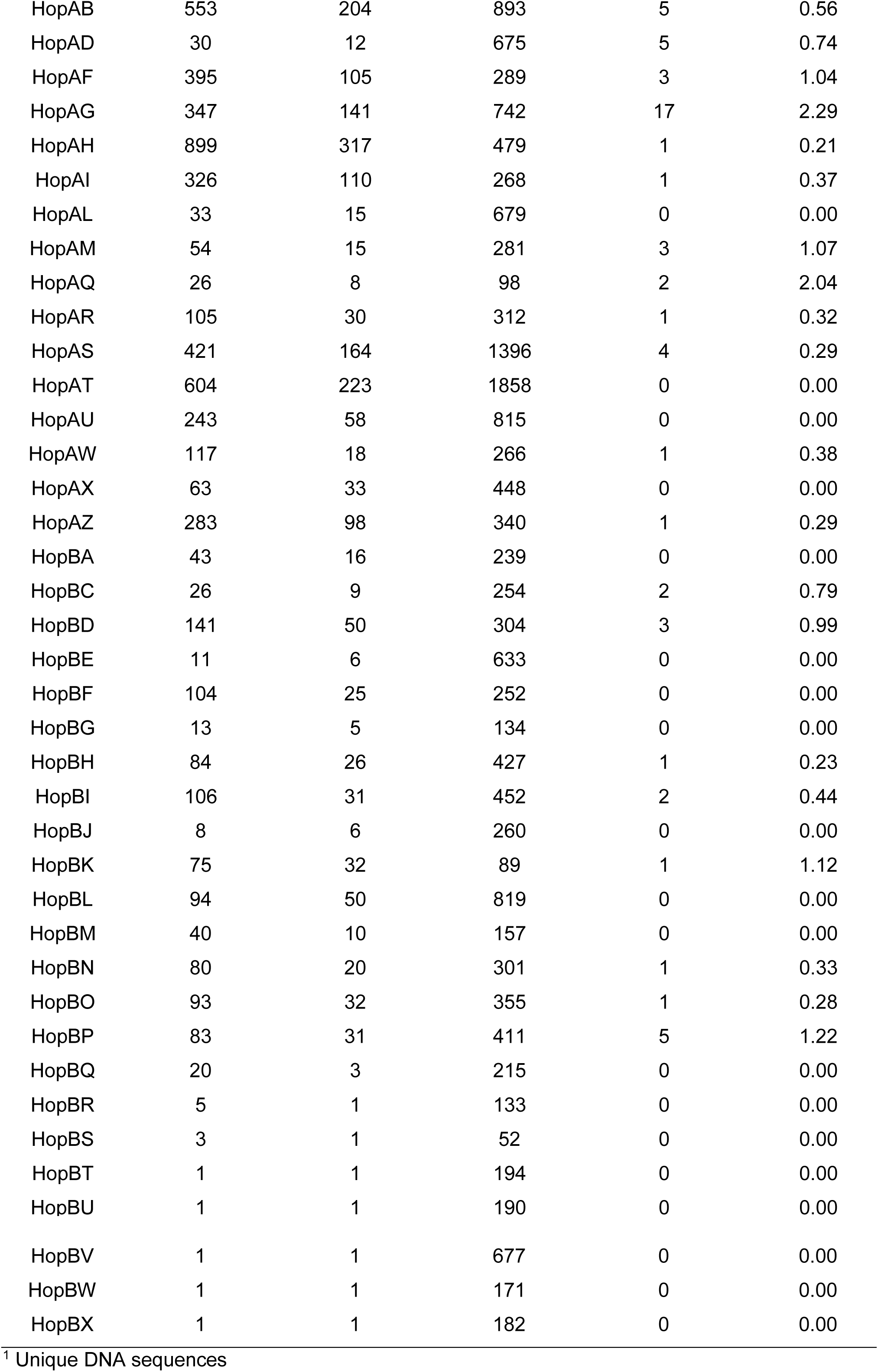
Positive Selection among T3SE Families.

| Family | Total Number of Alleles | Number of Unique Alleles <sup>1</sup> | Alignment Length (Codons) | Positively Selected Sites (N) | Positively Selected Sites (%) |
| --- | --- | --- | --- | --- | --- |
| AvrA | 27 | 12 | 906 | 1 | 0.11 |
| AvrB | 277 | 75 | 366 | 0 | 0.00 |
| AvrE | 608 | 360 | 2248 | 3 | 0.13 |
| AvrPto | 170 | 33 | 275 | 0 | 0.00 |
| AvrRpm | 171 | 39 | 301 | 0 | 0.00 |
| AvrRpt | 25 | 12 | 261 | 3 | 1.15 |
| HopA | 277 | 105 | 449 | 0 | 0.00 |
| HopB | 770 | 362 | 2265 | 0 | 0.00 |
| HopC | 115 | 28 | 271 | 0 | 0.00 |
| HopD | 587 | 228 | 981 | 0 | 0.00 |
| HopE | 103 | 31 | 274 | 0 | 0.00 |
| HopF | 380 | 125 | 385 | 0 | 0.00 |
| HopG | 190 | 70 | 528 | 0 | 0.00 |
| HopH | 265 | 54 | 226 | 2 | 0.88 |
| HopI | 400 | 166 | 601 | 0 | 0.00 |
| HopK | 156 | 34 | 338 | 3 | 0.89 |
| HopL | 102 | 53 | 902 | 1 | 0.11 |
| HopM | 620 | 223 | 1034 | 2 | 0.19 |
| HopN | 74 | 25 | 350 | 0 | 0.00 |
| HopO | 227 | 75 | 391 | 1 | 0.26 |
| HopQ | 304 | 86 | 504 | 3 | 0.60 |
| HopR | 424 | 231 | 2001 | 6 | 0.30 |
| HopS | 114 | 26 | 179 | 2 | 1.12 |
| HopT | 97 | 34 | 398 | 2 | 0.50 |
| HopU | 15 | 4 | 264 | 0 | 0.00 |
| HopV | 307 | 74 | 738 | 2 | 0.27 |
| HopW | 618 | 219 | 1125 | 1 | 0.09 |
| HopX | 308 | 83 | 452 | 3 | 0.66 |
| HopY | 201 | 53 | 287 | 2 | 0.70 |
| HopZ | 396 | 79 | 771 | 2 | 0.26 |
| HopAA | 752 | 218 | 578 | 0 | 0.00 |
| HopAB | 553 | 204 | 893 | 5 | 0.56 |
| HopAD | 30 | 12 | 675 | 5 | 0.74 |
| HopAF | 395 | 105 | 289 | 3 | 1.04 |
| HopAG | 347 | 141 | 742 | 17 | 2.29 |
| HopAH | 899 | 317 | 479 | 1 | 0.21 |
| HopAI | 326 | 110 | 268 | 1 | 0.37 |
| HopAL | 33 | 15 | 679 | 0 | 0.00 |
| HopAM | 54 | 15 | 281 | 3 | 1.07 |
| HopAQ | 26 | 8 | 98 | 2 | 2.04 |
| HopAR | 105 | 30 | 312 | 1 | 0.32 |
| HopAS | 421 | 164 | 1396 | 4 | 0.29 |
| HopAT | 604 | 223 | 1858 | 0 | 0.00 |
| HopAU | 243 | 58 | 815 | 0 | 0.00 |
| HopAW | 117 | 18 | 266 | 1 | 0.38 |
| HopAX | 63 | 33 | 448 | 0 | 0.00 |
| HopAZ | 283 | 98 | 340 | 1 | 0.29 |
| HopBA | 43 | 16 | 239 | 0 | 0.00 |
| HopBC | 26 | 9 | 254 | 2 | 0.79 |
| HopBD | 141 | 50 | 304 | 3 | 0.99 |
| HopBE | 11 | 6 | 633 | 0 | 0.00 |
| HopBF | 104 | 25 | 252 | 0 | 0.00 |
| HopBG | 13 | 5 | 134 | 0 | 0.00 |
| HopBH | 84 | 26 | 427 | 1 | 0.23 |
| HopBI | 106 | 31 | 452 | 2 | 0.44 |
| HopBJ | 8 | 6 | 260 | 0 | 0.00 |
| HopBK | 75 | 32 | 89 | 1 | 1.12 |
| HopBL | 94 | 50 | 819 | 0 | 0.00 |
| HopBM | 40 | 10 | 157 | 0 | 0.00 |
| HopBN | 80 | 20 | 301 | 1 | 0.33 |
| HopBO | 93 | 32 | 355 | 1 | 0.28 |
| HopBP | 83 | 31 | 411 | 5 | 1.22 |
| HopBQ | 20 | 3 | 215 | 0 | 0.00 |
| HopBR | 5 | 1 | 133 | 0 | 0.00 |
| HopBS | 3 | 1 | 52 | 0 | 0.00 |
| HopBT | 1 | 1 | 194 | 0 | 0.00 |
| HopBU | 1 | 1 | 190 | 0 | 0.00 |
| HopBV | 1 | 1 | 677 | 0 | 0.00 |
| HopBW | 1 | 1 | 171 | 0 | 0.00 |
| HopBX | 1 | 1 | 182 | 0 | 0.00 |

<sup>1</sup> Unique DNA sequences

Finally, to explore whether T3SE families display different levels of diversity than core gene families carried by the same *P. syringae* strains, we compared all pairwise *Ka* and *Ks* values within each effector family to the pairwise *Ka* and *Ks* values for the core genes carried in the corresponding genomes. We would expect T3SEs and core genes to share the same *Ka* and *Ks* values if they were evolving under the same evolutionary pressures. Deviation from this null expectation could be due to either differences in selective pressures, or the movement of the T3SE via horizontal gene transfer (HGT). We find that the pairwise *Ka* values for T3SEs are substantially higher than those of the corresponding core genes for the majority of T3SEs (Figure5A; Figure S7). This was also true for pairwise *Ks* values, although the differences between T3SE pairs and core genes were not as high and there were many more examples of T3SE pairs that had lower *Ks* values than the corresponding core genes (Figure 6B; Figure S8).

**Figure 6:**
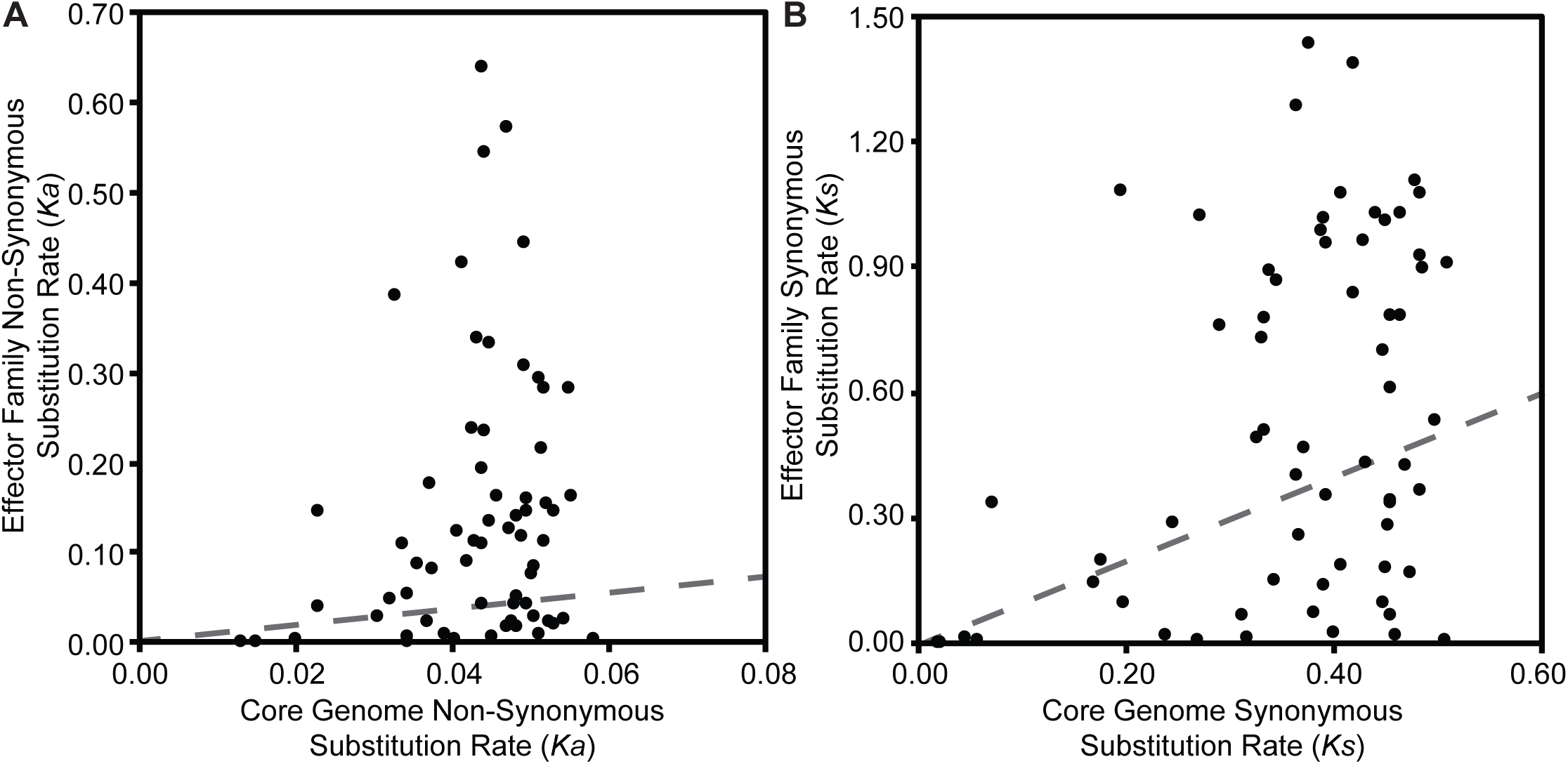
Relationship between the average pairwise non-synonymous substitution rate (*Ka*) (A) and the average pairwise synonymous substitution rate (*Ks*) (B) for each effector family with the average core genome synonymous and non-synonymous substitution rates of the corresponding *P. syringae* strains. Pairwise substitution rates for all sequences within a family were estimated by reverse translating the effector family and concatenated core genome amino acid alignments, then calculating pairwise substitution rates in MEGA7 with the Nei-Gojobori Method. Each point on the scatter plot represents the average of these pairwise rates for a single family and the red dotted lines represent the null-hypothesis that the substitution rates in the effector family will be the same as the substitution rates of the core genes in the same collection of genomes.

### Gene gain and loss of type III secreted effectors in the *P. syringae* species complex

Both the patchy distribution of T3SE families across the *P. syringae* species complex and the inconsistent relationships between T3SE and core gene substitution rates suggest that HGT may be an important evolutionary force contributing to the evolution of T3SEs in the *P. syringae* species complex. Therefore, we also sought to analyze the expected number of gene gain events across the *P. syringae* phylogenetic tree in order to more accurately quantify the extent to which HGT has actively transferred T3SEs between *P. syringae* strains over the evolutionary history of the species complex. We used the Gain Loss Mapping Engine (GLOOME) to estimate the number of gain and loss events (Cohen et al., 2010;Cohen and Pupko, 2010), and found extensive evidence for HGT in several T3SE families, with some families experiencing as many as 40 HGT events over the course of the history of the *P. syringae* species complex (Figure 7). Outlier T3SE families that did not appear to have undergone much HGT in *P. syringae* include the smallest families, like HopU, HopBE, and HopBR, and the largest families, like AvrE, HopB, HopM, and HopAA. Smaller families were less likely to have undergone HGT because they were only identified in a subset of closely related strains, so are not expected to have been part of the *P. syringae* species complex through the majority of its evolutionary history. Larger families may experience less HGT because they are more likely to already be present in the recipient strain and therefore will quickly be lost following an HGT event. However, because GLOOME only identifies HGT events that result in the gain of a new family, we cannot be certain whether *P. syringae* genomes with multiple copies were generated by HGT or gene duplication.

**Figure 7:**
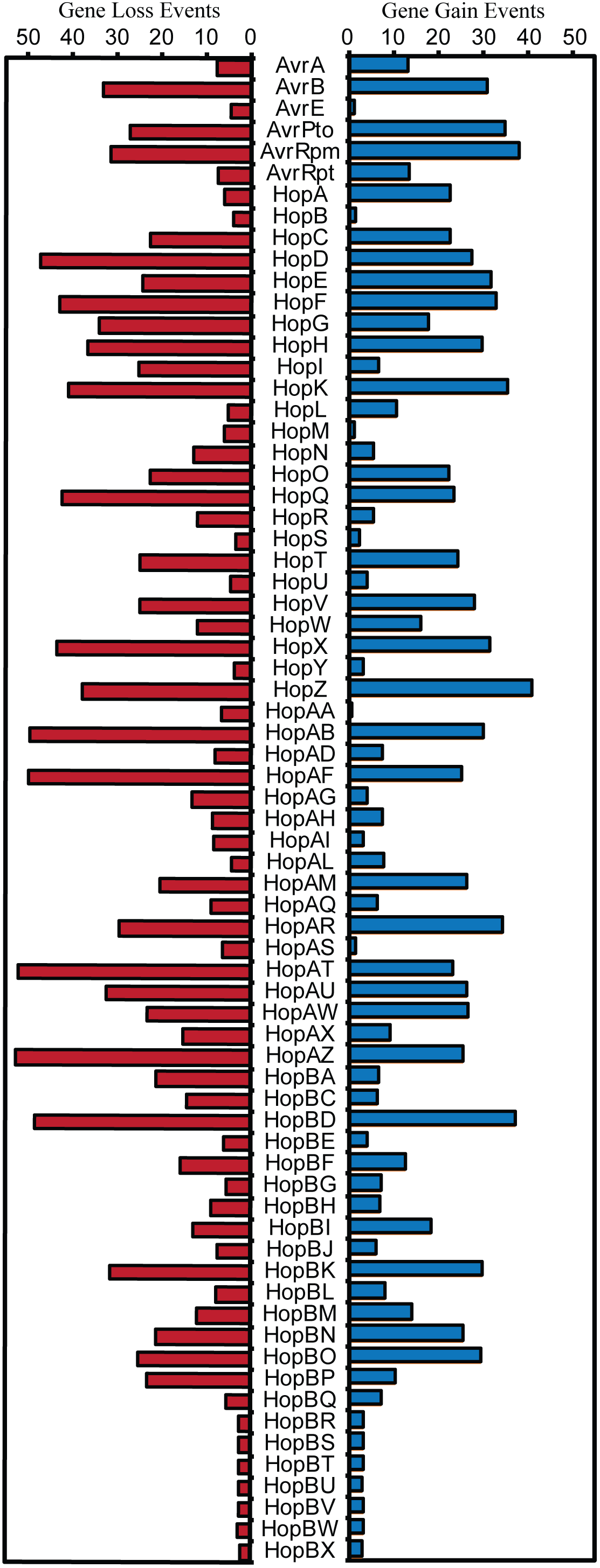
Expected number of gene gain and gene loss events for each T3SE family. The posterior expectation for gain and loss events was estimated for each family on each branch of the *P. syringae* core-genome tree using GLOOME with the stochastic mapping approach. The sum of these posterior expectations across all branches yields the total expected number of events for each family.

An opposing evolutionary force that is also expected to have a disproportional effect on the evolution of T3SE families is gene loss. Specifically, loss of a given T3SE may allow a *P. syringae* strain to infect a new host by shedding an effector that elicits the hosts’ ETI response. Indeed, we found that gene loss events were also common in many T3SE families, with more than 50 events estimated to have occurred in the HopAT and HopAZ families (Figure 7). T3SE families that experienced more gene loss events also tended to experience more gene gain events, as demonstrated by a strong positive correlation between gene loss and gene gain in T3SE families (Figure S9) (linear regression; F = 140.50, df = 1, 68, p < 0.0001, r^2^ = 0.67). However, as was the case with gene gain events, we observed few gene loses in the smallest and the largest T3SE families. For small families, this is again likely to be the result of the fact that they have spent less evolutionary time in the *P. syringae* species complex. For large families, we are again blind to gene loss events that occur in a genome that has multiple copies of the effector prior to the loss event. Therefore, there are likely many more T3SE losses occurring in larger families than we observe here because these T3SE families tend to be present in multiple copies within the same genome.

Finally, we also observed that there is a significant positive correlation between both evolutionary rate parameters and the rates of gene gain and loss for T3SE families (*Ka*-Gene Gain: F = 8.48, df = 1, 63, p = 0.0050, r^2^ = 0.1186; *Ka*-Gene Loss: F = 16.15, df = 1, 63, p = 0.0002, r^2^ = 0.2041; *Ks*-Gene Gain: F = 6.46, df = 1, 63, p = 0.0135, r^2^ = 0.0930; Ks-Gene Loss: F = 7.70, df = 1, 63, p = 0.0072, r^2^ = 0.1089) (Figure S10). This implies that the same evolutionary forces resulting in diversification of T3SEs are also causing them to undergo elevated rates of gain or loss. However, there was substantial unexplained variance in these correlations, resulting in some T3SE families that have high evolutionary rates and low levels of gain and loss, and other T3SE families that have low evolutionary rates and high levels of gain and loss. These families tended to be the same for all correlations.

## DISCUSSION

Bacterial T3SEs are primary virulence factors in a wide-range of plant and animal pathogens (Hueck, 1998;Desveaux et al., 2006;Zhou and Chai, 2008;Block and Alfano, 2011;Buttner, 2016;Khan et al., 2016;Hu et al., 2017;Khan et al., 2018;Xin et al., 2018). T3SEs are particularly interesting from an evolutionary perspective due to their dual and diametrically opposed roles in host-pathogen interactions. While T3SEs have evolved in order to promote bacterial fitness, usually via the suppression of host immunity or disruption of host cellular homeostasis, hosts have evolved mechanisms to recognize the presence or activity of T3SEs, and this recognition often elicits an immune response that shifts the interaction back into the host’s favor. To explore the distribution and evolutionary history of *P. syringae* T3SEs and gain insight into their role in host specificity, we catalogued the T3SE repertoires of a large and diverse collection of 494 *P. syringae* isolates. These phylogenetically diverse strains allowed us to generate an expanded database of more than 14,000 T3SE alleles and investigate the evolutionary mechanisms through which these important molecules have enabled *P. syringae* to become one of the most globally important bacterial plant pathogens (Mansfield et al., 2012).

### Expanded database of type III secreted effectors in *P. syringae*

This study increases the number of confirmed and putative T3SE alleles available in the *P. syringae* Genome Resources Database by 20-fold, resulting in a final database of 14,613 T3SE alleles from the *P. syringae* species complex, 5,127 of which are unique at the nucleotide level. Although these new, putative T3SEs all share an ancestral sequence with known T3SE families, the extensive diversification that has occurred within many of these families clearly indicates that some level of functional diversification has occurred.

Consistent with our earlier analysis, we find that primary phylogroup strains harbor considerably larger repertoires of T3SEs than secondary phylogroup strains (Baltrus et al., 2011;O’Brien et al., 2011;Dudnik and Dudler, 2014;Dillon et al., 2017). We also find that a small number of primary phylogroup strains have significantly smaller effector repertoires; including phylogroup 10 strains, which were primarily isolated from non-agricultural sources similar to most secondary phylogroup strains, and the phylogroup 2 strain Psy642, which has previously been highlighted as an outlier in its T3SE content and has been characterized as non-pathogenic (Clarke et al., 2010;O’Brien et al., 2011). In general, phylogroup 2 strains have somewhat smaller T3SE repertoires and employ a greater number of phytotoxins relative to other primary phylogroup strains (Baltrus et al., 2011;O’Brien et al., 2011; Dillon et al., 2017). This may indicate that phylogroup 2 strains have evolved a different host-microbe lifestyle than other *P. syringae* primary phylogroup strains, e.g. one tending towards low virulence, epiphytic interactions, rather than high virulence, invasive pathogenesis (Hirano and Upper, 2000).

Among the 70 T3SE families that were delimited in this study, seven of them had fewer than five total members (HopBR, HopBS, HopBT, HopBU, HopBV, HopBW, HopBX). These families all consist of alleles that were separated from a larger T3SE family during the delimitation stage of our analysis because they shared only very limited regions of local similarity with the larger family. The small size of these families suggests that they may be pseudogenes degenerating due to a lack of selective constraints. The 63 remaining families are similar to the ~60 families that have been discussed in earlier studies (Baltrus et al., 2011;Lindeberg et al., 2012). While we do merge seven families based on our delimitation analysis, seven new families have been discovered in the past five years (McCann et al., 2013;Hockett et al., 2014;Lam et al., 2014;Matas et al., 2014;Mucyn et al., 2014). Unfortunately, our objective delimitation analysis separated HopX2 from HopX, HopZ3 from HopZ, and HopH3 from HopH, forming the HopBO, HopBP, and HopBQ families, respectively. Despite these differences, we arrive at several similar conclusions to prior work on the distribution of individual T3SEs across *P. syringae* strains. Specifically, we find that few T3SE families are considered part of the core genome (Baltrus et al., 2011;O’Brien et al., 2011;Lindeberg et al., 2012), with only AvrE, HopB, HopM, and HopAA being present in more than 95% of strains. Three of these families (AvrE, HopM, and HopAA) are part of the CEL, while the other CEL effector, HopN, is only present in 14.98% of *P. syringae* strains, all from phylogroup 2. This suggests that HopN arose in the CEL after the divergence of this phylogroup. Other families that have previously been characterized as core T3SEs in *P. syringae* include HopI and HopAH (Baltrus et al., 2011), which are only present in 79.76% and 89.07% of strains from our study, respectively. HopB has not been highlighted as a core T3SE in prior studies, likely because it had been split into the HopB and HopAC families. We find that alleles from these families are quite similar, often sharing reciprocal BLASTP hits across more than 80% of the HopB sequence with E-values less than 1e-24, which indicates that HopB and HopAC should be considered a single family. The remainder of T3SE families have a considerably sparser distribution across the *P. syringae* species complex, ranging in frequency from 1.62% to 80.97%. This demonstrates that different T3SE families were likely acquired episodically throughout the evolutionary history of the *P. syringae* species complex and are subject to strong evolutionary pressures for gain and loss due to the widespread and diverse ETI surveillance system of plants (Cunnac et al., 2009;Xin et al., 2018).

Finally, we find that highly divergent combinations of T3SEs can enable *P. syringae* to infect the same host (Figure S6). While this observation is consistent with prior studies in *P. syringae* (Baltrus et al., 2011;Lindeberg et al., 2012;O’Brien et al., 2012), it is in contrast to the convergence in T3SE repertoires that has been observed in *Xanthomonas*, another phytopathogen that employs a T3SS (Hajri et al., 2009). Importantly, this limits our ability to detect and differentiate *P. syringae* pathogens of different hosts using this fairly crude application of comparative genomics. The lack of correlation between T3SE repertoires and host specificity may be a direct result of the fact that there is substantial functional redundancy among *P. syringae* T3SEs from different families, or that certain T3SEs in combination can mask the detection of other T3SEs in a given *P. syringae* background (Cunnac et al., 2009;Cunnac et al., 2011;Lindeberg et al., 2012;Wei et al., 2018). However, it will be important moving forward to assess the true host range of a broader collection of *P. syringae* strains in order to determine whether specific T3SEs promote or suppress growth on particular hosts.

### Genetic and functional evolution of *P. syringae* type III secreted effectors

Given the broad array of unique T3SEs that exist within the *P. syringae* species complex, mining this untapped diversity is likely to reveal a number of new functions and interactions for T3SEs in *P. syringae*. By quantifying *Ka, Ks*, and *Ka/Ks* for each pair of T3SE alleles in each family, we identified substantial genetic diversity in several T3SE families (Figure 5). Our codon-level analysis of positive selection also revealed that T3SE families were substantially more likely than non-T3SE families to contain positively selected sites (Table 3). Finally, we confirmed that this divergence is not simply a reflection of the immense diversity exhibited by the strains used in this study, since the divergence observed for T3SE families is consistently higher than the divergence observed across core genes (Figure S7; Figure S8). Elevated non-synonymous substitution rates in T3SE families implies that there is elevated positive selection operating on these families. Elevated synonymous substitution rates additionally show that this elevated positive selection may extend to synonymous sites, that many T3SEs arose prior to the last common ancestor (LCA) of the *P. syringae* species complex, and/or that T3SEs undergo considerably higher rates of HGT than core genes.

Fast-evolving T3SEs will also provide numerous opportunities for studying Red Queen dynamics (van Valen, 1973). Under Fluctuating Red Queen (FRQ) dynamics, fluctuating selection drives oscillations in allele frequencies at the focal genetic loci in both the pathogen and the host, resulting in rapid evolutionary change on both sides (Brockhurst et al., 2014). In the case of *P. syringae* and their plant hosts, bacterial T3SEs are the key players on the pathogen side, and plant resistance genes are the key players on the host side. These FQR dynamics are expected to maintain high levels of within-population genetic diversity at focal loci, as we’ve observed in many T3SE families. The majority of T3SE families in *P. syringae* are highly divergent and display strong signatures of positive selection, likely in response to intense host-imposed selection to evade recognition (Rohmer et al., 2004;Baltrus et al., 2011;Lindeberg et al., 2012). This implies that few T3SEs are broadly unrecognized, making interactions between individual T3SEs and the corresponding plant resistance genes an excellent resource for exploring FQR dynamics.

The highly dynamic nature of T3SE evolution is also seen in our analysis of T3SE gain and loss across the *P. syringae* phylogenetic tree. More than five gene gain events are estimated to have occurred in 52 out of the 70 T3SE families analyzed in this study, with a maximum of 41 HGT events estimated in the HopZ family. Gene loss events were even more common, with 57 out of 70 T3SE families experiencing more than five loss events and a maximum of 53 events in the HopAZ family. Earlier studies have also suggested that both gene gain and loss were quite common among T3SE families. One specific study using nucleotide composition and phylogenetics found that members from 11 out of 24 tested *P. syringae* T3SE families were recently acquired by HGT (Rohmer et al., 2004). These families included AvrA, AvrB, AvrD, AvrRpm, HopG, HopQ, HopX, HopZ, HopAB, HopAF, and HopAM (although AvrD is not a T3SE (Leach and White, 1996;Mucyn et al., 2014)). The T3SEs from this dataset were also highlighted by this study as undergoing considerably high rates of gene gain and loss within the *P. syringae* species complex. Specifically, all of these T3SEs were demonstrated to have undergone at least ten gene gain events and many were among the most dynamic T3SEs in our dataset. Other studies have shown that many T3SEs are present on mobile genetic elements and that T3SEs from the same family are often found at different genomic locations (Kim and Alfano, 2002;Charity et al., 2003;Lovell et al., 2009;Godfrey et al., 2011;Lovell et al., 2011;Neale et al., 2016), which may both promote and be a consequence of the high rates of gene gain and loss for particular T3SE families. From a selective perspective, it is also likely that host immune recognition can drive selection for gene gain or loss (Vinatzer et al., 2006), while the functional redundancy of different T3SE families carried in the same genetic background may limit the negative impacts of the loss of such T3SEs (Kvitko et al., 2009;Cunnac et al., 2011;Wei et al., 2018). Finally, as has been previously reported (Baltrus et al., 2011), we find that there is a significant positive correlation between rates of evolution and rates of gene gain and loss (Figure S10), suggesting that similar evolutionary forces that cause the diversification of T3SEs are contributing to the loss and gain of T3SEs. However, not all T3SEs fit this model which could reflect that T3SEs vary in their mutational robustness and/or that the genomic context of different T3SEs makes them more or less prone to HGT. In any event, the extensive gene gain and loss that occurs in the majority of T3SE families lends further support to the hypothesis that few T3SE alleles are broadly unrecognized (Baltrus et al., 2011).

Given the highly dynamic nature of T3SE evolution, we predict that there are still numerous T3SEs that will be found to elicit ETI. Most research on ETI elicitation to date has focused on a small number of T3SE families, and an even smaller number of alleles from each family (Mansfield, 2009). The immense diversification that we observe in many T3SE families points to strong selective pressures that may be explained by as-yet discovered ETI responses. If this prediction holds true, it will be particularly interesting to study T3SE families with alleles that induce different ETI responses in the same host. These patterns will help reveal how strains shift onto new hosts or break immunity in an existing host, perhaps explaining the evolutionary driving force behind new disease outbreaks.

## DATA ACCESS

All genomic data produced by this study have been submitted to NCBI. Accession numbers for all genomes sequenced in this study and all publicly available genomes are available in Supplemental Dataset S1.

## ACKNOWLEDGMENTS

We thank all members of the Guttman and Desveaux labs for helpful discussion and valuable input on this project. This work was supported by Natural Sciences and Engineering Research Council of Canada Discovery Grants (D.S.G and D.D.), Canada Research Chairs in Comparative Genomics (D.S.G.) and Plant-Microbe Systems Biology (D.D.), and the Center for the Analysis of Genome Evolution and Function (D.S.G. and D.D.).

## AUTHOR CONTRIBUTIONS

M.M.D., D.D., and D.S.G. designed the research; M.M.D., R.A., B.L., and A.M. analyzed the data; and M.M.D, and D.S.G. wrote the paper.

