## Supplemental Material for "Molecular Evolution of *Pseudomonas syringae* Type III Secreted Effector Proteins"

**Table of Contents**

1. **SUPPLEMENTAL FIGURES	3**
   1.1. Supplemental Figure S1	3
   1.2. Supplemental Figure S2	4
   1.3. Supplemental Figure S3	5
   1.4. Supplemental Figure S4	6
   1.5. Supplemental Figure S5.	7
   1.6. Supplemental Figure S6	8
   1.7. Supplemental Figure S7	9
   1.8. Supplemental Figure S8	10

1.10. Supplemental Figure S9	11

1.11. Supplemental Figure S10	12

2. **SUPPLEMENTAL DATASETS	13**
   2.1. Supplemental Dataset S1	13
   2.2. Supplemental Dataset S2	14

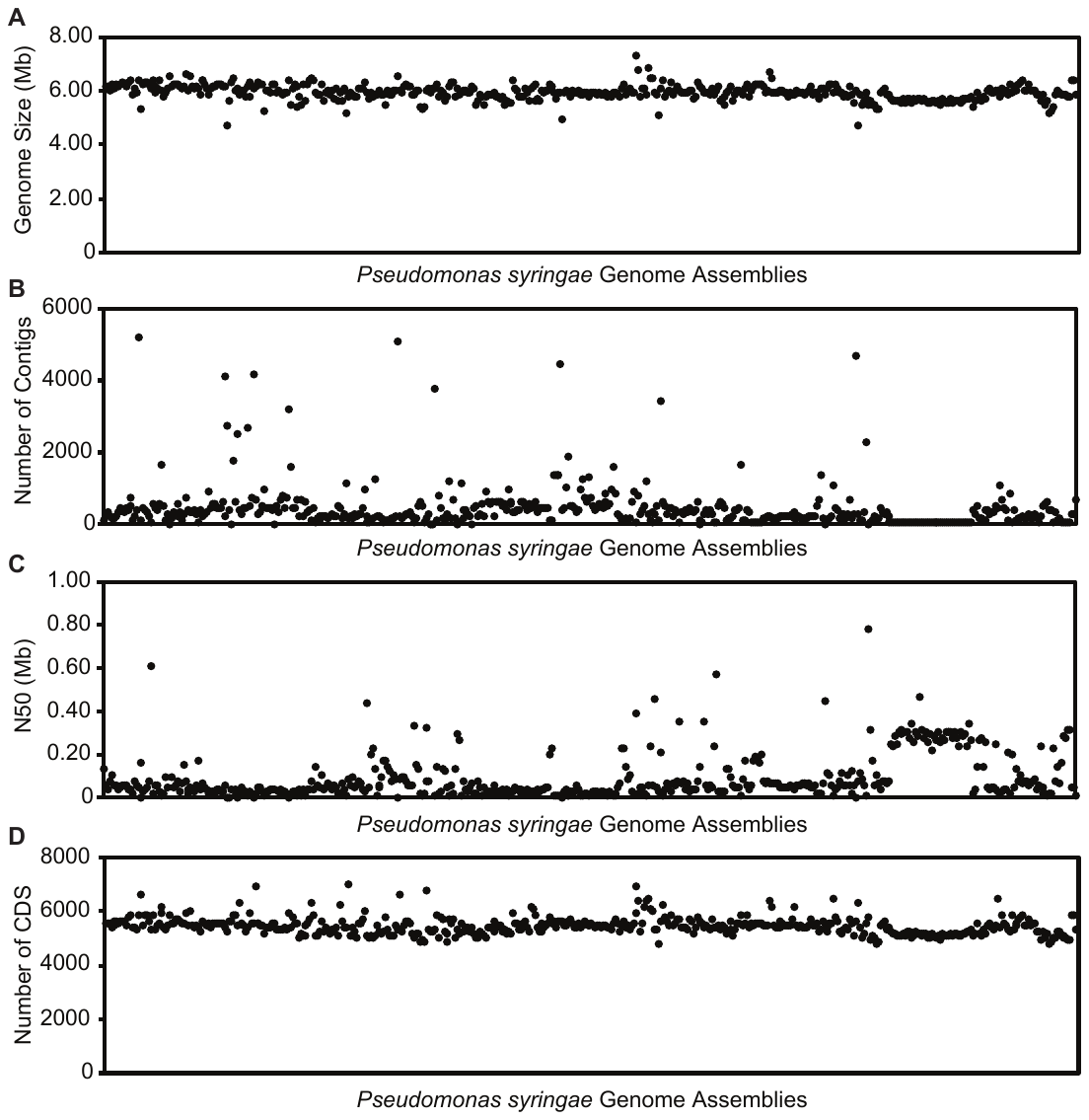

**Figure S1: Assembly summary for all *Pseudomonas syringae* genomes used in this study.** The genome sizes (A), number of contigs (B), N50 values (C), and number of coding sequences (D) of all the *P. syringae* assemblies.

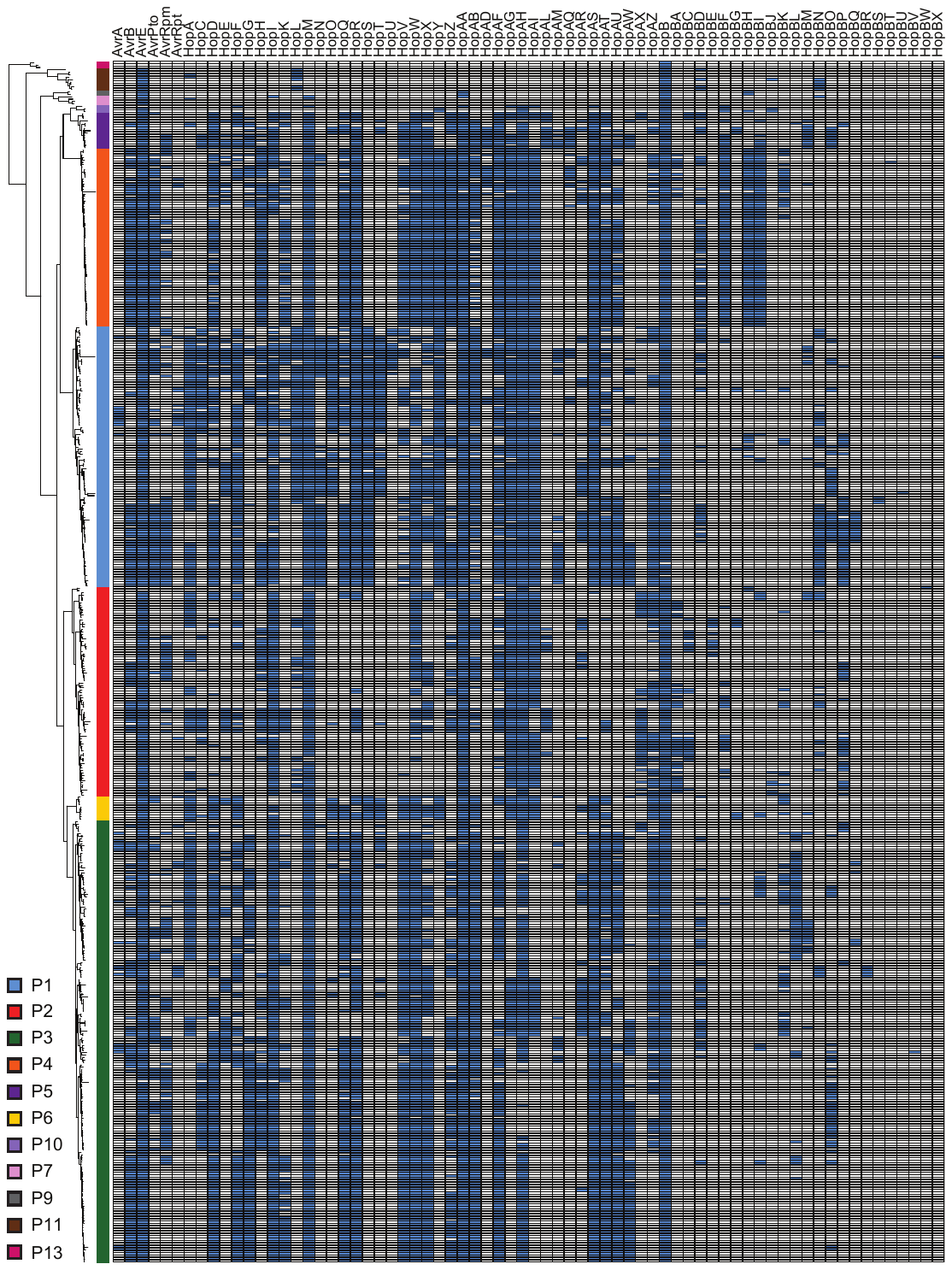

**Figure S2: Distribution of type III secreted effectors across the *Pseudomonas syringae* species complex.** Filled boxes represent significant blast hits for the effector family in an annotated coding sequence of each strain, as described in the Methods.

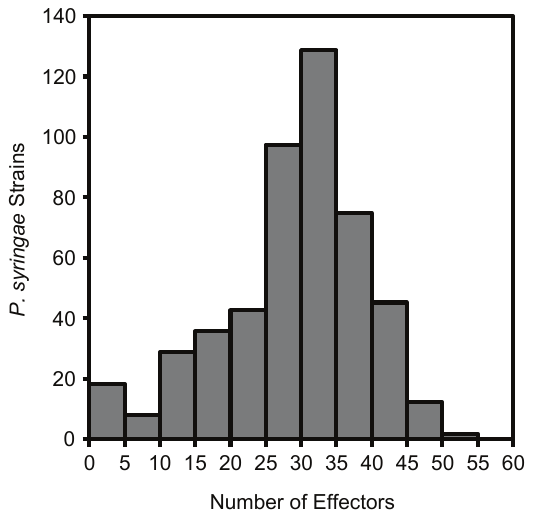

**Figure S3: Distribution of the number of type III secreted effectors in each of the 494 *Pseudomonas syringae* species used in this study**.

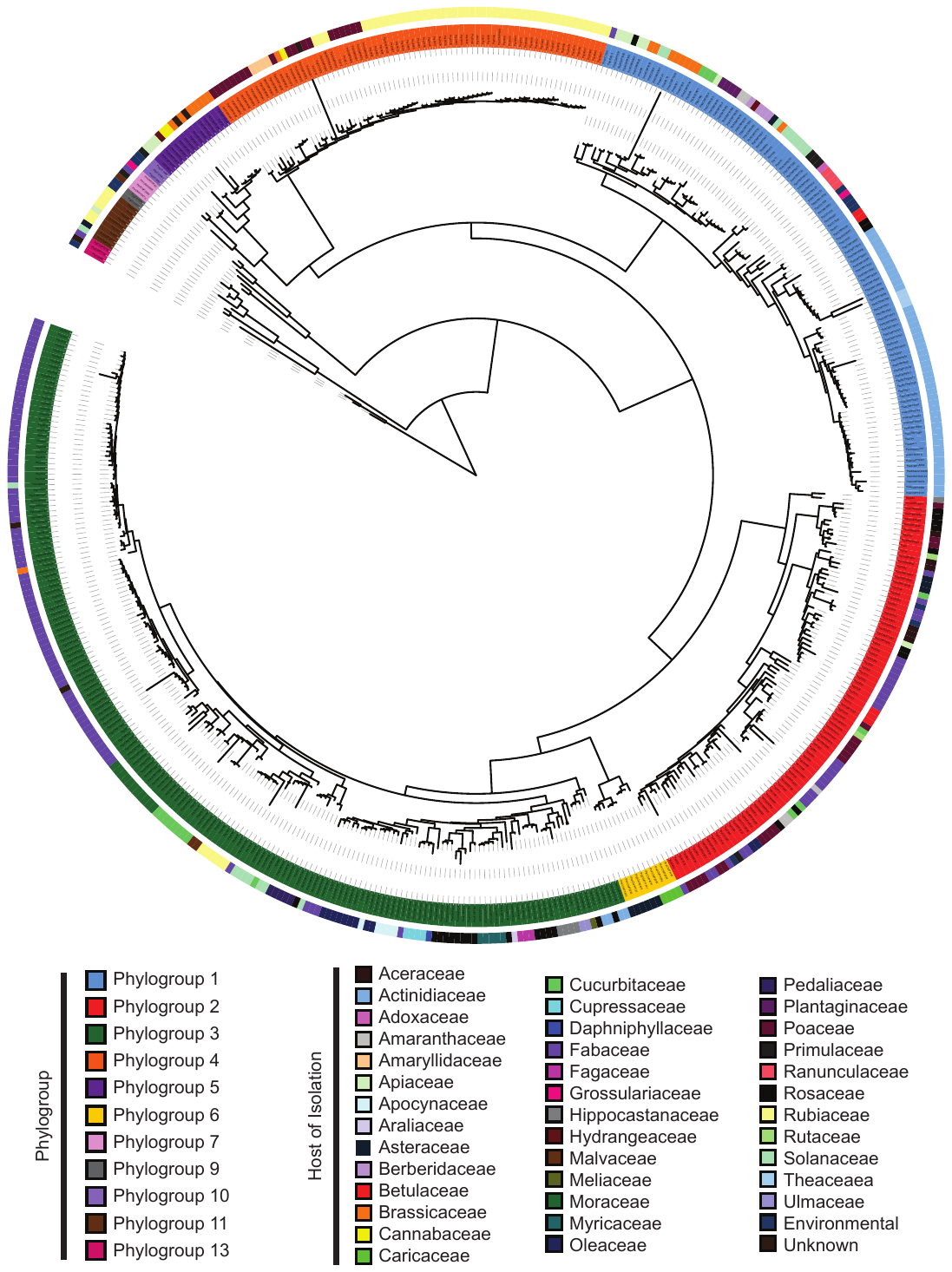

**Figure S4: Core genome phylogenetic tree of all *Pseudomonas syringae* strains used in this study.** The core genome maximum-likelihood tree was generated from a core genome alignment of the 1,324 core genes of the *P. syringae* strains analyzed in this study. Strain phylogroups and host families are shown outside the tree.

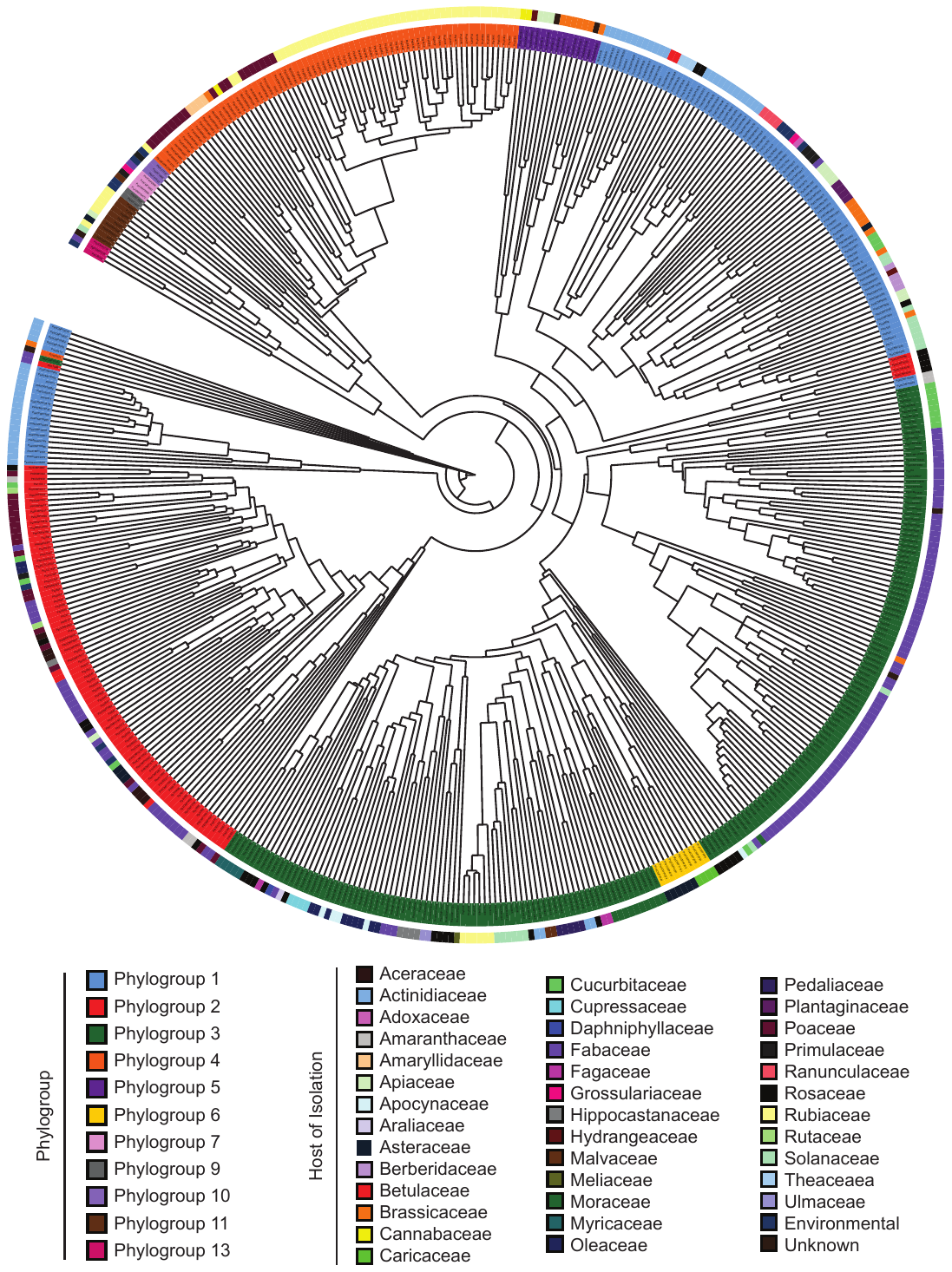

**Figure S5: Pan-genome phylogenetic tree of all *Pseudomonas syringae* strains used in this study.** The pan-genome tree was generated by hierarchical clustering of the gene content in each strain. Strain phylogroups and host families are shown outside the tree.

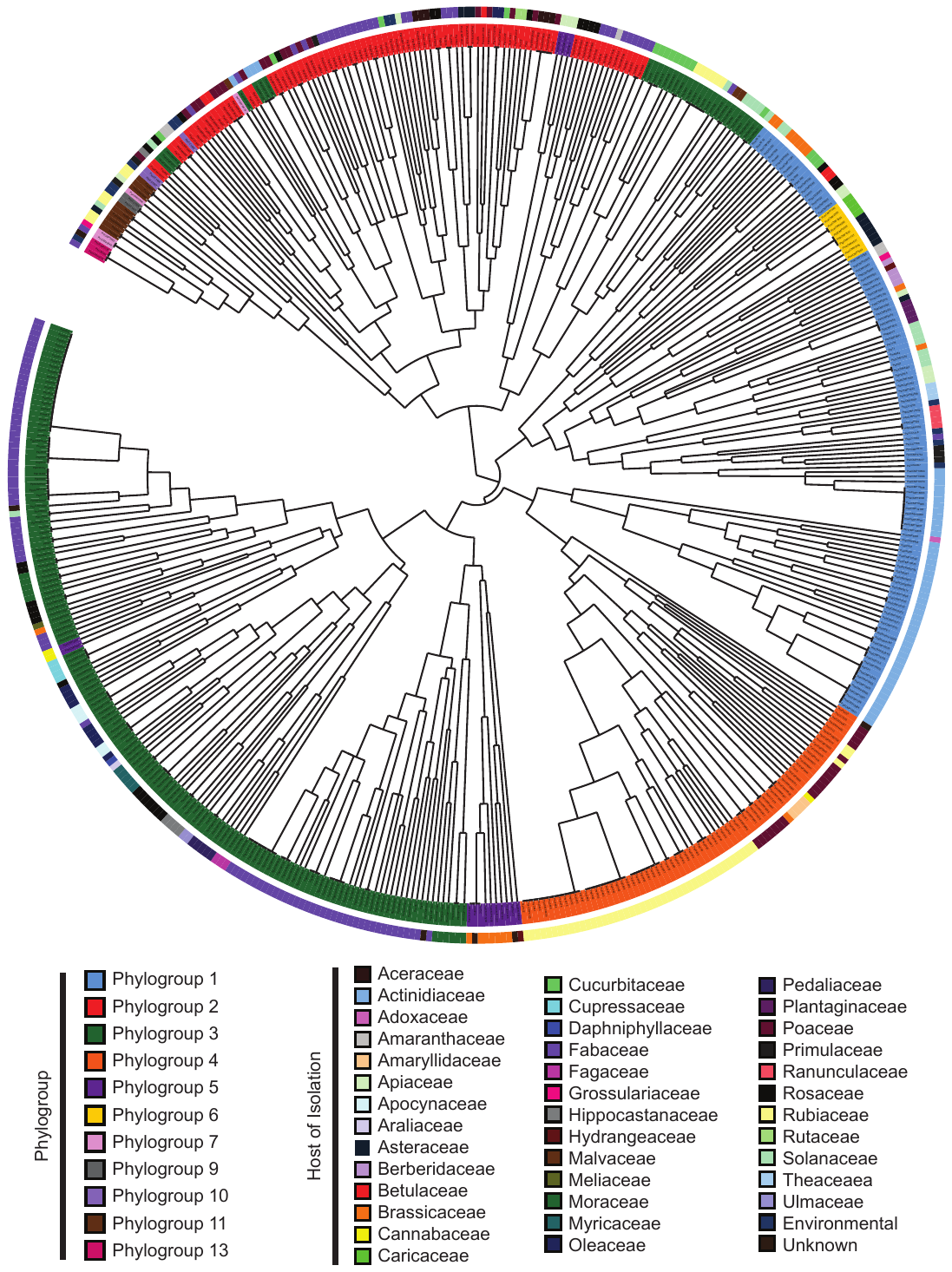

**Figure S6: Effecterome phylogenetic tree of all *Pseudomonas syringae* strains used in this study.** The effecterome tree was generated by hierarchical clustering of the effector content in each strain. Strain phylogroups and host families are shown outside the tree.

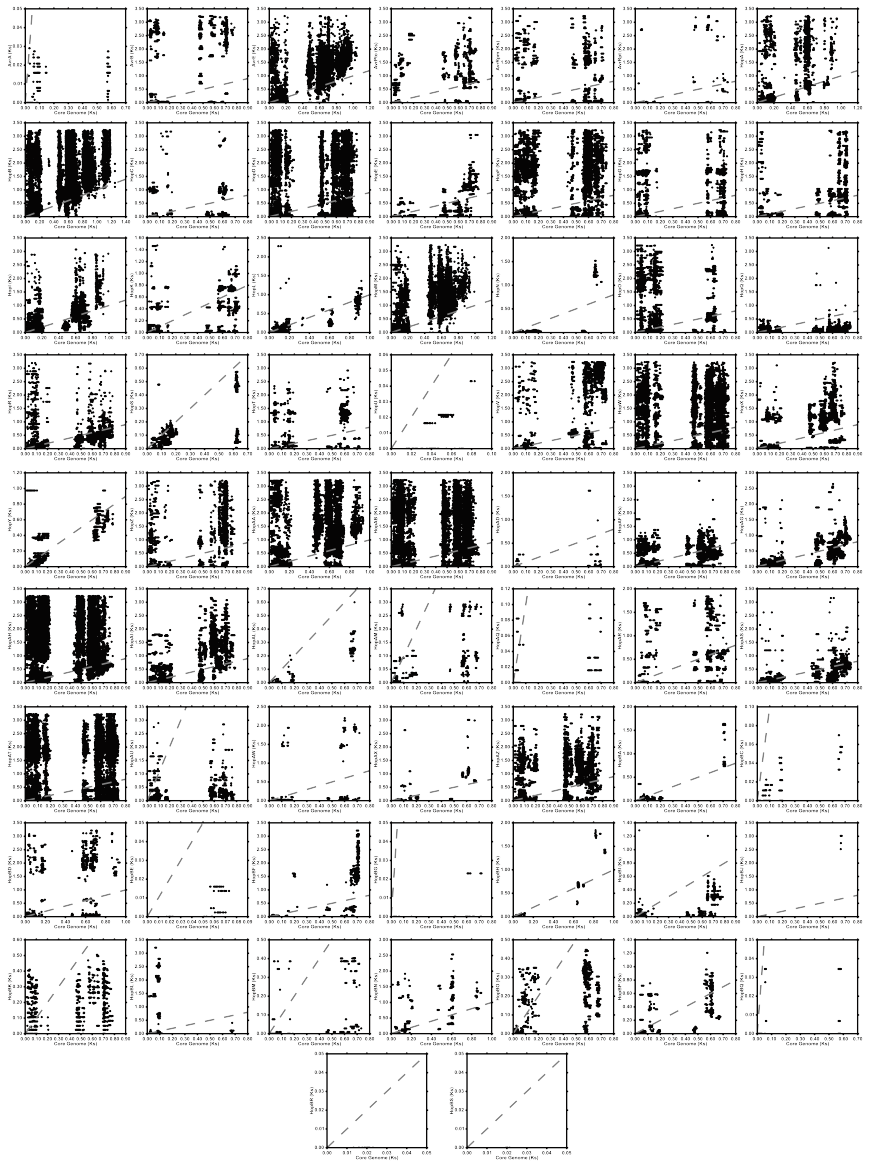

**Figure S7: Relationship between all pairwise synonymous substitution rates (Ks) within each type III effector family and corresponding core genome synonymous substitution rates (Ks) of the appropriate strain pairs.** Pairwise substitution rates were estimated by reverse translating the concatenated core genome and effector family amino acid alignments to nucleotide alignments, then calculating pairwise synonymous substitution rates in MEGA7 with the Jukes-Cantor Method. Grey dotted lines represent the null-hypothesis that the pairwise synonymous substitution rate of the effector allele pair will be equal to the rates estimated from the core genomes of those strains.

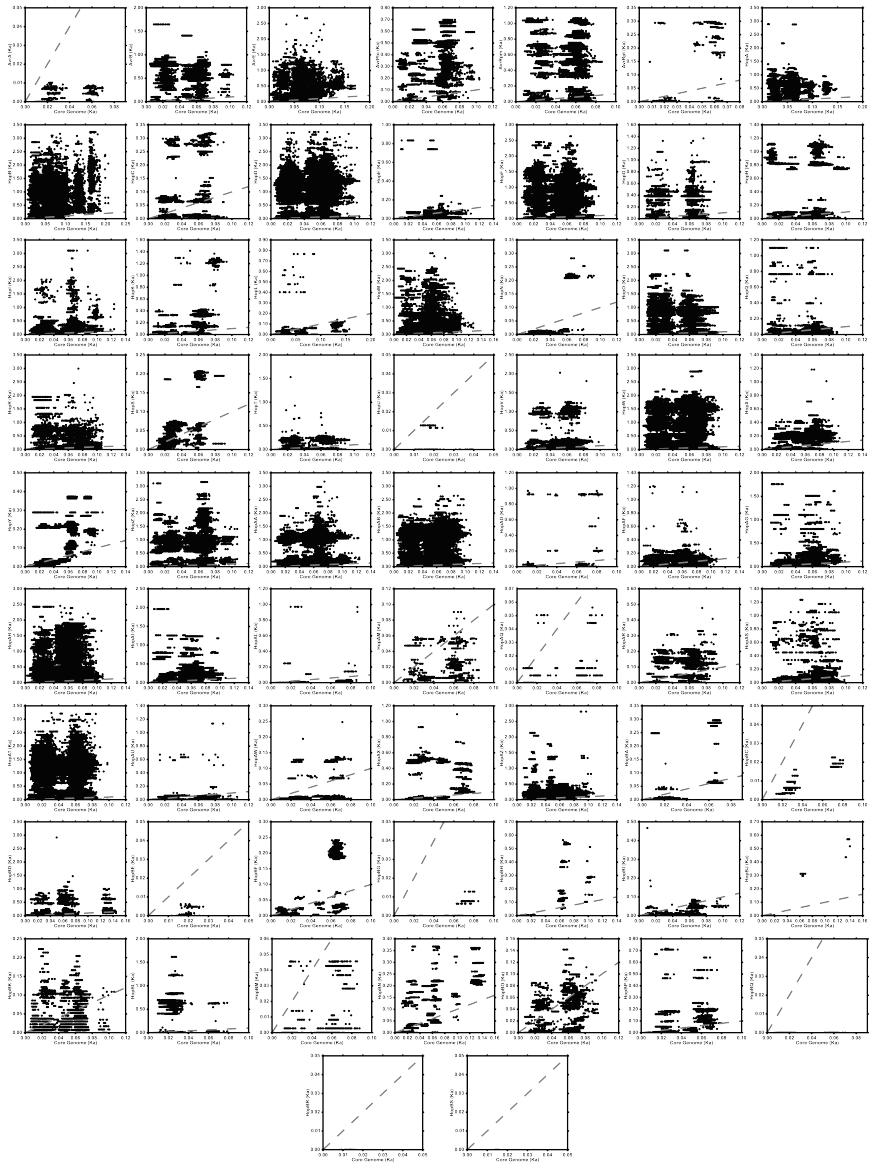

**Figure S8: Relationship between all pairwise non-synonymous substitution rates (Ka) within each type III effector family and corresponding core genome non-synonymous substitution rates (Ka) of the appropriate strain pairs.** Pairwise substitution rates were estimated by reverse translating the concatenated core genome and effector family amino acid alignments to nucleotide alignments, then calculating pairwise synonymous substitution rates in MEGA7 with the Jukes-Cantor Method. Grey dotted lines represent the null-hypothesis that the pairwise non-synonymous substitution rate of the effector allele pair will be equal to the rate estimated from the core genomes of the corresponding strains.

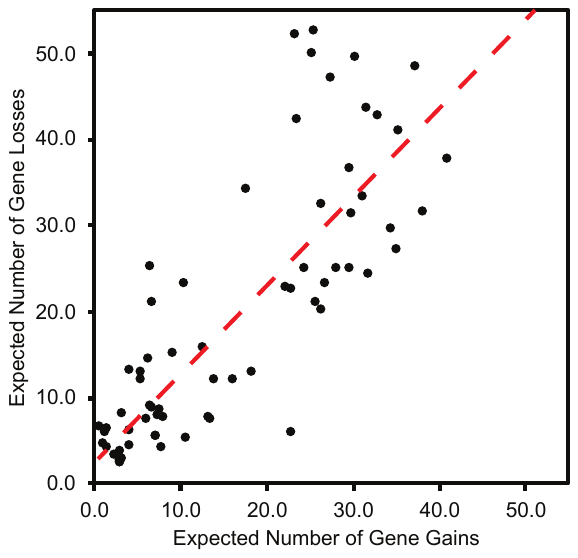

**Figure S9: Relationship between the expected number of gene gain events and the expected number of gene loss events estimated to have occurred for each type III effector family over the evolutionary history of the *P. syringae* species complex (F = 140.50, df = 1, 68, p < 0.0001, r^2^ = 0.6691).** For each family, the posterior expectation of gene gain and loss events were estimated for each branch of the phylogenetic tree using the stochastic mapping method implemented with GLOOME. The sum of these events across all branches yields the expected number of events per family. The linear regression is illustrated by the red dashed line.

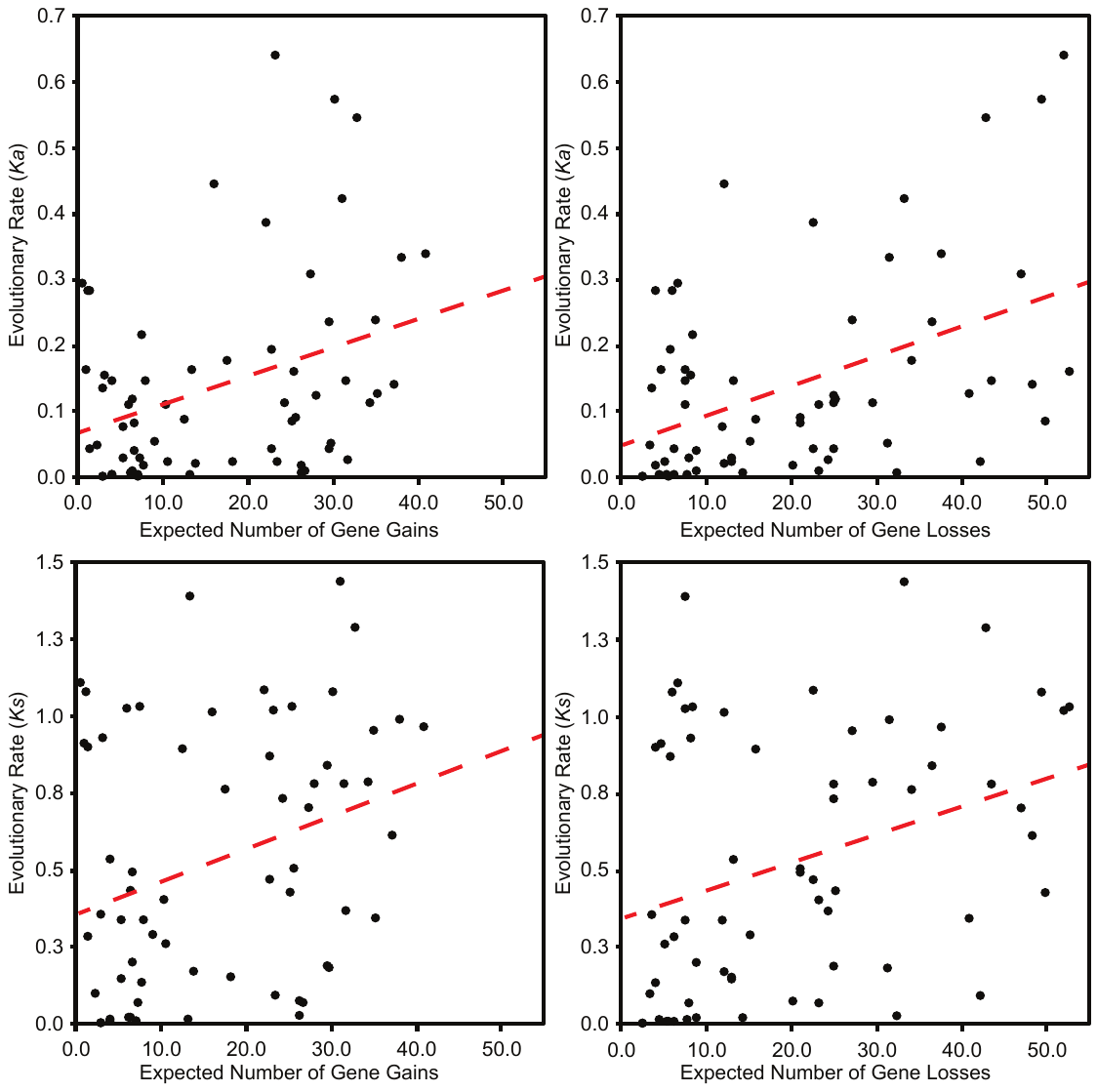

**Figure S10: Relationship between the expected number of gene gain/loss events and evolutionary rate parameters, *Ka* and *Ks*, for each type III effector family over the evolutionary history of the *P. syringae* species complex.** A) Relationship between the average non-synonymous substitution rate in each T3SE family and the expected number of gene gain events (F = 8.48, df = 1, 63, p = 0.0050, r^2^ = 0.1186). B) Relationship between the average non-synonymous substitution rate in each T3SE family and the expected number of gene loss events (F = 16.15, df = 1, 63, p = 0.0002, r^2^ = 0.2041). C) Relationship between the average synonymous substitution rate in each T3SE family and the expected number of gene gain events (F = 6.46, df = 1, 63, p = 0.0135, r^2^ = 0.0930). D) Relationship between the average synonymous substitution rate in each T3SE family and the expected number of gene loss events (F = 7.70, df = 1, 63, p = 0.0072, r^2^ = 0.1089). Linear regressions are illustrated by the red dashed lines in each panel.

**Supplemental_Dataset_S1: Strain information for all 494 *P. syringae* species complex genomes used in this study.**

Supplemental_Dataset_S1.xlsx

**Supplemental_Dataset_S2: Summary information for all type III secreted effectors identified in this study and their family designations.**

Supplemental_Dataset_S2.xlsx
